# Integrated Continuous Biomanufacturing of Recombinant Adeno-Associated Virus

**DOI:** 10.64898/2026.09.14.751635

**Authors:** Richard Plieninger, Julia M. Müller, Daniela Tobler, Efe Saygili, Patrick Werder, Yuki Higuchi, Ryosuke Takahashi, Sebastian Vogg, Thomas Müller-Späth, Sven Göbel, Thomas K. Villiger

## Abstract

Recombinant adeno-associated virus (rAAV) manufacturing remains constrained by low yields, costly purification, and process complexity, limiting cost-effective access to gene therapy. Here, we demonstrate an integrated continuous biomanufacturing platform that transfers proven antibody-manufacturing technology to rAAV production by coupling perfusion to semi-continuous twin-column affinity capture (CaptureSMB). This architecture enables continuous harvest of extracellular rAAV without cell lysis, depth filtration, or endonuclease treatment, sustaining stable operation over five days. Relative to conventional batch processing, perfusion improved capsid or vector genome yield depending on the production process, while affinity capture alone reduced total DNA to batch-comparable levels, with no measurable loss in impurity clearance or potency. By consolidating multiple manual unit operations into one continuous workflow, the platform may reduce manual intervention and process footprint. Its modular design may generalize to other rAAV serotypes, inducible producer cell lines, and viral vector classes, offering a potential route toward more efficient gene therapy manufacturing.

**Graphical abstract:** 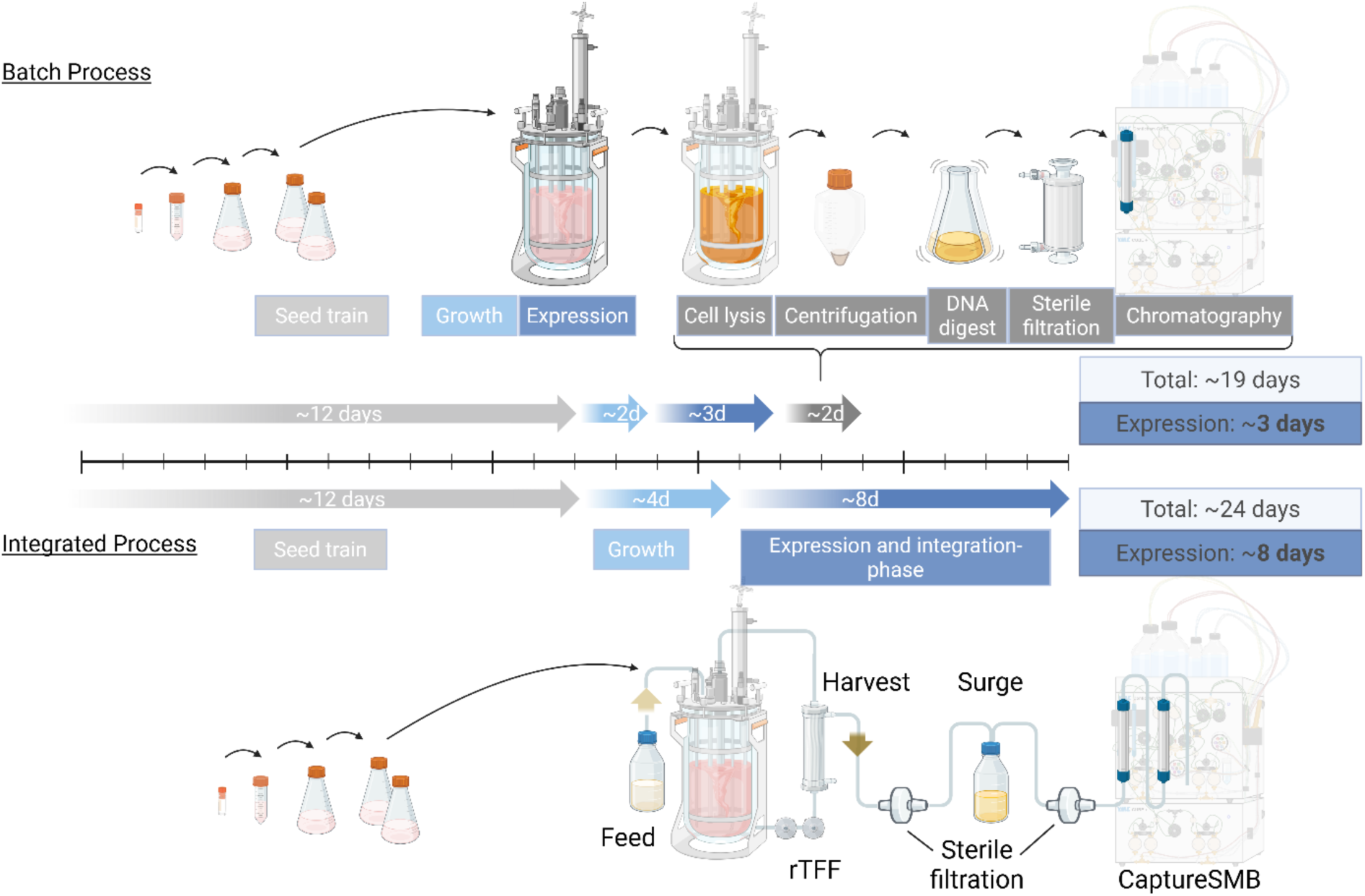

**Highlights:**

- An integrated continuous biomanufacturing platform for rAAV by coupling perfusion to a twin-column CaptureSMB affinity system has been established.
- Bypassed traditional, costly, and yield-limiting batch operations including chemical cell lysis, depth filtration, and endonuclease treatment by capturing exclusively extracellular rAAVs.
- Transfer of various scalable continuous process technologies from antibody processes to rAAV manufacturing.
- Alternative platform with the potential to increase overall yields, lower cost of goods, increase automation and sustain or even improve critical quality attributes.

**Technology readiness:** Whilst individual continuous processing units such as perfusion bioreactors and multi-column chromatography systems are mature and already employed in industry for various biologics, continuous production and purification systems for recombinant adeno-associated virus (rAAV) are still in their infancy. Recent studies suggest intensifying rAAV workflows via upstream perfusion and multi-column chromatography, however the integration of these steps into a cohesive continuous manufacturing line has remained a conceptual-stage challenge Technology Readiness Level (TRL 2). In this manuscript, we have established a proof-of-concept at laboratory scale conditions where state-of-the-art technologies from integrated continuous biomanufacturing for antibodies have been leveraged and applied to rAAV biomanufacturing. Thus, all employed technological elements used are available with scalable options for industrial implementation. Consequently, integrated continuous biomanufacturing of rAAV has advanced through TRL 3 (experimental proof of concept) to TRL 4 (technology validated in the laboratory) of the integrated continuous biomanufacturing workflow. In order to progress to TRL 5 (technology validated in relevant environment), it is necessary to demonstrate higher and consistent process yield, robust product quality, scalability and operational practicality under manufacturing conditions.

## Introduction

Recombinant adeno-associated viruses (rAAVs) have become the predominant viral vector platform for in vivo gene therapy, with programs in over 700 pre- and 225 clinical stages and eight FDA-approved products by 2026 [1,2]. While low-dose therapies (e.g., Luxturna, 1.5 × 10^11^ vector genomes (vg) per eye [3]) can utilize established adherent HEK systems [4], systemic administrations require higher vector doses, typically ranging from 1 × 10^14^ to 1 × 10^16^ vg per patient [1]. Most modern platforms rely on suspension cells to bridge this dose-manufacturing gap, yet they face substantial obstacles. Large-scale rAAV manufacturing is challenged by low production yields, limited inherent scalability [5], and the need for rigorous optimization to ensure high full/empty capsid ratios and minimize process-related impurities like host cell DNA and host cell proteins (HCPs). As a result, the estimated cost-of-goods (COGs) for a single gene therapy treatment (1 x 10^15^ vg/dose) ranges from US$ 12,000 –100,000 per dose, underscoring the urgent need for improved manufacturing efficiencies to expand rAAV access beyond ultra-rare indications [6].

In response to prevailing manufacturing constraints and the imperative to reduce COGs, continuous manufacturing has emerged as a transformative paradigm in bioprocessing [7]. The shift from sequential batch processing to a constant, simultaneous flow of material across interconnected steps has demonstrated significant COGs reductions in monoclonal antibody (mAb) manufacturing: up to 68% for clinical and 35% for commercial scales [8]. Continuous processing offers further advantages including reduced downtime, smaller footprint, higher equipment utilization, increased overall titer, decreased processing steps and higher product quality, mostly due to lower product residence time that is critical for sensitive products such as viral vectors [9,10]. While mAb processes benefit from optimized stable cell lines and mature process development, the comparatively young field of rAAV gene therapy lacks an explicitly standardized production process. Nevertheless, the core unit operations are generally analogous to those in mAb manufacturing, making continuous processing technologies developed for antibody processes highly adaptable to viral vector manufacturing, in particular perfusion and multi-column chromatography.

Perfusion represents an effective process intensification strategy for cell culture-based recombinant protein manufacturing. Filter-based cell retention technologies, notably tangential flow filtration (TFF) and alternating tangential flow filtration (ATF), are the most prevalent systems in industrial bioprocessing, enabling precise nutrient delivery, efficient metabolite removal, and continuous product harvest for direct downstream capture, thereby maximizing volumetric productivity and overall process efficiency [11,12]. Recent advancements have addressed sieving-related limitations in filter-based perfusion systems through the incorporation of a secondary pump in a reverse-TFF (rTFF) configuration and alternating activation, coupled with targeted optimization of filter length and fiber diameter [13–15]. These modifications effectively reduce the impact of Starling recirculation and thus increase filter performance. These factors are critical for prolonged high cell concentration perfusion for efficient rAAV harvesting, given the large size of viral particles and accumulation of host cell-derived debris. Advantages of perfusion for rAAV production include not only the drastic increase of cell concentration prior to transfection [16] but also having the possibility of dilution after high cell density (HCD) transfection [17] and the option for final filtration at the end of the culture [16]. Hence, perfusion processes have been developed for various viral vector constructs including modified vaccinia Ankara virus [18] and lentivirus production [9].

For rAAVs, continuous extracellular rAAV harvesting has been demonstrated for multiple serotypes, including rAAV2, rAAV5, rAAV8 [16,19,20]. However, extracellular yields three days post-transfection are strongly serotype-dependent, ranging from 0.5–11% for rAAV2, 12–28.1% for rAAV5, and 19–42.1% for rAAV8 [21,22]. Moreover, continuous secretion of rAAV5 has been shown for up to 19 days post-transfection increasing the cumulative yield over 4-fold compared to traditional recovery from cellular lysates [19]. Nevertheless, for most rAAV serotypes such as rAAV2, a significant proportion of rAAV particles remain intracellular especially for shorter expression times like the classical 2-3 days post-transfection for batch runs, and require cell lysis for release during harvest [21]. This lysis step drastically increases the release of process-related impurities (HCP and total DNA), significantly burdening downstream processing (DSP). Consequently, the addition of endonucleases such as benzonase is necessary to reduce long-stranded DNA load prior to filtration and chromatographic purification, a step that can account for up to 51% of total upstream material costs [23]. Most workflows then proceed with depth-filtration (or centrifugation at smaller scales) and 0.2 µm filtration. Depth-filtration often results in substantial rAAV losses, with reported yields ranging from 60-90% [24].

Similarly, continuous DSP has also advanced, with proposals for continuous counter-current multicolumn capture schemes using two, three or four-column processes in combination with different affinity columns [25–27]. Furthermore, implementation of a multicolumn countercurrent solvent gradient purification (MCSGP) strategy has demonstrated superior efficacy in achieving filled particle enrichment when compared to traditional batch purification methodologies [28,29]. Although these progresses confirm that continuous manufacturing approaches from other biologics can be transferred to rAAV, an integrated continuous biomanufacturing system for rAAV production has not been reported to the authors’ knowledge.

The purpose of this study is the integration of state-of-the art perfusion technology and continuous capture process to an rAAV process. More precisely, reverse tangential flow filtration (rTFF) perfusion is coupled directly to twin-column CaptureSMB. This architecture enables the direct capture of extracellular rAAVs, bypassing the traditional requirements for lysis, endonuclease treatment, primary clarification and up concentration prior to affinity capture. We evaluated the performance of this platform for two distinct rAAV5 production processes, providing a comparison of each integrated continuous process against its respective batch process.

## Material and methods

### Cell culture system

HEK293F cells (Viral Production Cells 2.0, Thermo Fisher, Waltham, MA, USA) were grown in HyClone Peak Expression Medium (Cytiva, Marlborough, MA, USA) and cultivated in an incubator (Multitron, Infors HT, Bottmingen, Switzerland) at 36.5 °C and 5% CO2. The orbital throw for 50 mL spin tubes (Tube Spin, TPP, Trasadingen, Switzerland) with 10 - 30 mL cell culture was set to 50 mm at 190 rpm, otherwise 25 mm at 140 rpm for bigger cell expansion in flasks. For each transient transfection three plasmids were used. For Process A: pHelper (pJD171 #229483), pRepCap (pAAV2/5 #104964) and pTransfer (70_pAAV-ProA23-CatCh-GFP-WPRE #125906 [30] or pAAV.CMV.PI.EGFP.WPRE.bGH #105530) and for Process B: pHelper (pADΔF6 #112867), pRepCap (pAAV2/5 #104964), and pTransfer (pAAV.CMV.PI.EGFP.WPRE.bGH #105530), obtained from Addgene, Watertown, MA, USA and amplified and purified in-house [31]. For the transient transfection, 2 μg of total plasmid DNA per million cells was mixed in a molar ratio of 1:2:0.5 (pTransfer: pRepCap: pHelper), and subsequently complexed using polyethylenimine Max (PEI Max, Polysciences, Warrington, PA, USA) in a 2:1 ratio, complexed in medium (10% of the culture volume) for 15 min before adding to the cells.

### Batch process

Cells were transferred to bioreactors (Labfors 5, Infors HT) with a seeding concentration of 1 × 10^6^ cells/mL with a working volume of 1.6 L. Bioreactors were equipped with pH (EasyFerm Plus Arc, Hamilton, Bonaduz Switzerland), optical DO (VisiFerm DO Arc, Hamilton), biocapacitance (Incyte Arc, Hamilton), and pCO_2_ (CO_2_NTROL RS485, Hamilton) probes. After reaching a cell concentration of 2 × 10^6^ cells/mL, cells were transfected. Dissolved oxygen level was controlled by a cascade at 50%, temperature at 36.5 °C, the flat disc stirrer was set at 270 rpm. Samples were taken daily indicated in Figure 1 A as *c_Reactor_*. Cells were harvested 72 h post-transfection and directly lysed in the bioreactor using 0.5% (v/v) Tween-20 (P1379, Sigma-Aldrich International, Buchs, Switzerland), at 37 °C with agitation at 270 rpm for 1 h (Figure 1 A as *c_Lysis_*). After adjusting the pH to 7.4 using 0.2 M NaOH and final concentration of 150 mM NaCl (S9625, Sigma-Aldrich International), cell lysate was clarified by centrifugation (1000 g for 30 min, followed by 1500 g for 30 min) (Figure 1 A as *c_Centrif._*). Subsequently, DNA was digested using the endonuclease benzonase (Merck, 1016560001) at 50 U/mL and 2 mM MgCl₂ (M0250, Sigma-Aldrich International) for 2 h at 37 °C and 115 rpm (Figure 1 A as *c_Endo._*). To stabilize the rAAV particles, NaCl (S9625, Sigma-Aldrich International) was added to final 150 mM NaCl. Lastly, 0.2 μm bioburden reduction filtration was performed (Mini Kleenpak Capsule, KM5EAVP2S, Cytiva) (Figure 1 A as *c_SF_*).

**Figure 1.**
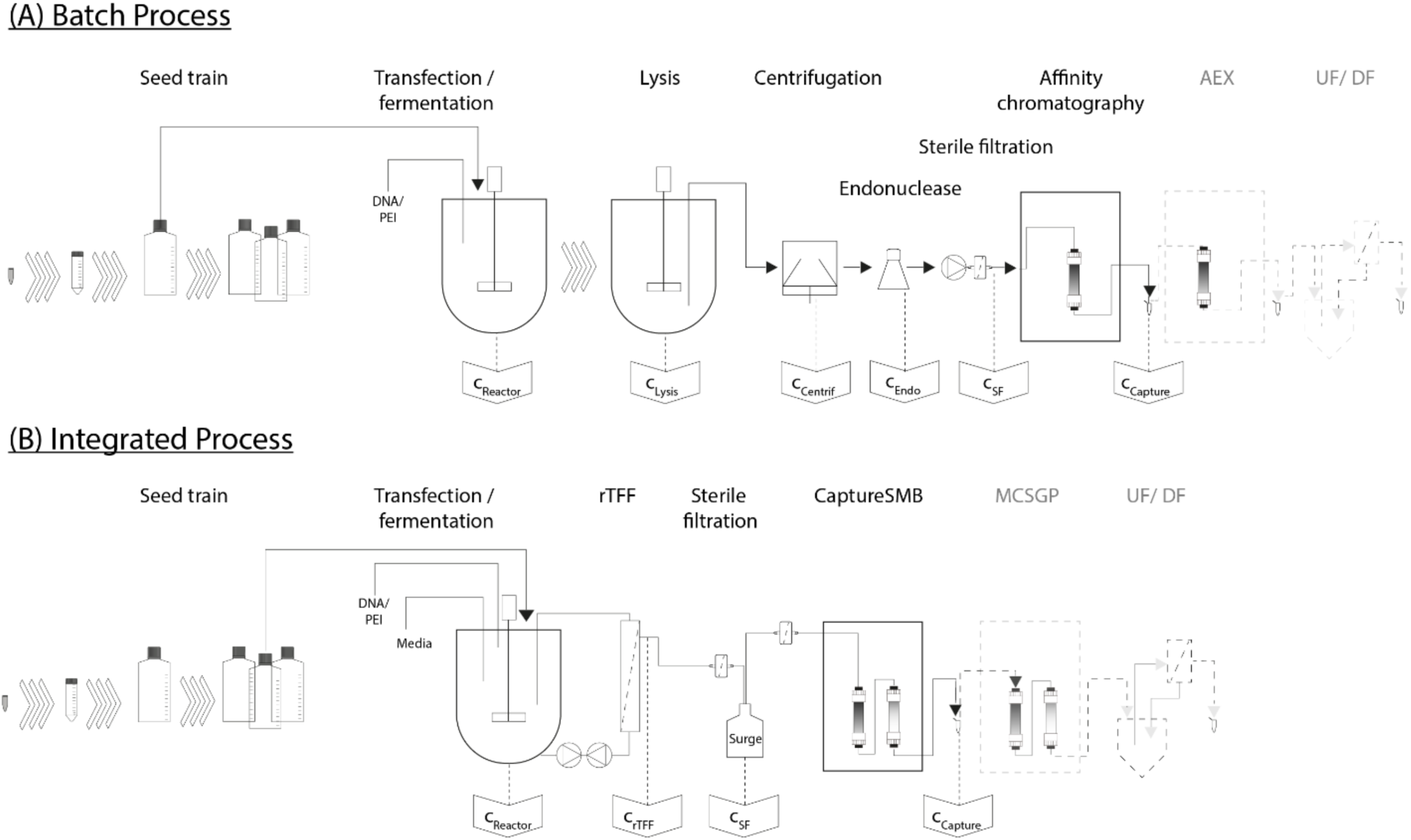
Process flow diagram of batch and integrated continuous rAAV production. (A) Batch process based on triple-plasmid transient transfection, followed by batch lysis, centrifugation, filtration and batch capture as well as polishing with final UF/DF. (B) Integrated continuous biomanufacturing approach, with a perfusion bioreactor connected to a surge tank, sterile filtration and twin-column capture chromatography. For both schemes, the final polishing chromatography and UF/DF steps are represented by dashed lines to illustrate a complete production stream; these final steps were not evaluated in this study.

Preparative chromatography experiments were conducted on a Contichrom® CUBE 30 system (ChromaCon AG, Zurich, Switzerland) controlled via ChromIQ® version 9.0 software. Conductivity, pH, and UV absorbance (260 and 280 nm) were monitored online. At each column outlet, UV detection was performed using a 0.5 mm optical path length. rAAV5 capture was performed using AVIPureAAV5 affinity chromatography resin (0.5 mL, 5 × 25 mm; Repligen, Waltham MA, USA) as previously described [25]. Briefly, the column was equilibrated with 10 column volumes (CV) of Tris-buffered saline (pH 8.0). Subsequently, the clarified harvest was loaded to 1.1% of the dynamic binding capacity (DBC) at a residence time of 0.45 min (1.1 mL/min), followed by a post-load wash with equilibration buffer (10 CV). Elution (Figure 1 A as *c_Capture_*) was achieved using a low-pH glycine/arginine buffer (pH 2.5), and the eluate was immediately neutralized using Tris-HCl (pH 10.0). Finally, the column underwent regeneration, cleaning-in-place with NaOH, neutralization, and re-equilibration according to the established protocol.

### Integrated continuous process

Perfusion cultivations were conducted using the same bioreactor configuration and cascade control strategy as described for the batch runs, unless otherwise stated below. Upon reaching a viable cell concentration (VCC) of 3 × 10^6^ cells/mL, perfusion was initiated at 0.5 vessel volumes per day (vvd). Cell retention was achieved using a 0.19 m² hollow-fiber module (MF-SL, 0.4 µm; Asahi Kasei, Tokyo, Japan) connected to two centrifugal BPS-i30 pumps (Levitronix, Zürich, Switzerland). When the culture reached 5 × 10^6^ cells/mL, transfection was performed and pump operation was switched from TFF to rTFF mode (recirculation flow rate: 1.1 L/min; flow direction alternated every 60 s) to ensure high product sieving [14]. Simultaneously, the perfusion rate was increased to 1 vvd. Process integration was initiated at a targeted flow rate of 1.1 mL/min after three days post-transfection. The harvest stream was directed to an elevated 1 L surge tank positioned upstream of the chromatography device to enable proper liquid transfer for the process integration. Sterile filter capsules (0.2 µm Supor KA02EAVP8G; Cytiva) were installed both upstream and downstream of the surge tank to ensure sterility. The surge tank was gravimetrically monitored, and the working volume was maintained between 150-200 mL (9 –13% of the bioreactor volume).

The filtered surge tank material was continuously captured using a CaptureSMB process as previously described [25]. In this approach two 0.5 mL AVIPureAAV5 affinity columns were operated in a cyclic manner on the Contichrom^®^ CUBE 30 system, alternating between interconnected and batch configurations. The key modification in this study was the adjustment of the interconnected load duration and reduced flow rate during batch loading (0.313 to 0.937 mL/min). To ensure consistent loading across these cycles, these parameters were calculated using historical titer profiles and total vector particle loads from the reference perfusion processes, ensuring that each switch achieved the target load established in [25]. During integrated runs, 17 and 11 cycles were performed for Process A and B, respectively. An overview of the downstream process parameters between batch and continuous chromatography is provided in Table S1.

Samples were collected once to twice a day at the following locations during the run as indicated in Figure 1 B: 1) directly from the bioreactor, *c_Reactor_* with lysed samples and samples followed by centrifugation at 1000 × g for 3 min, with supernatant retained for analysis; 2) from the permeate line after the rTFF, *c_rTFF_;* 3) from the surge tank with previous passing sterile filtration (SF), c_SF_; and 4) pooled elution fractions obtained after CaptureSMB affinity capture, *c_Capture_*.

### Cell culture analytics

VCC, culture viability, cell diameter, metabolites (glucose, glutamine, glutamate, lactate, ammonium), and pH were measured using a BioProfile FLEX2 (Nova Biomedical, Waltham, MA, USA). Here, bioreactors were automatically sampled 2–4 times per day by the FLEX2 On-Line Autosampler (Nova Biomedical). Additional aliquots of 1 mL were taken for further analysis.

### Virus quantification

To determine the proportion of extracellular capsids, samples were immediately centrifuged after collection at 1000 × g for 5 min at room temperature and the supernatant was collected. For total capsid quantification in cell broth, samples were lysed with 10 x lysis buffer (200 mM Tris, 1.5 M NaCl, 20 mM MgCl_2_, 5% Polysorbate 20 at pH 7.3) and incubated for 2 h at 36 °C in a thermomixer (Thermomixer comfort, Eppendorf, Hamburg, Germany) at 600 rpm. Quantitative rAAV5 capsid detection was performed using the ELISA kit (AAV5 Xpress ELISA PROGEN, Heidelberg, Germany) according to the manufacturer’s instructions. Genome copy number of encapsulated rAAV5 was determined by qPCR targeting the EGFP region of the transgene. Primers and probe were adapted from [32]: forward 5’-GAA CCG CAT CGA GCT GAA-3, reverse 5’-TGC TTG TCG GCC ATG ATA TAG-3’, and TaqMan probe 5’-ATC GAC TTC AAG GAG GAC GGC AAC-3’ (5’ FAM, 3’ BHQ-1). Not-encapsulated DNA was removed via DNase digest. A total of 20 µL reaction was prepared including, 2 µL of sample, 13.2 µL DNase free water, 2 µL 10 × Pluronic F-68 solution, 0.8 µL DNase I (2 U/µL; M0303S NEB, Ipswich, MA, USA) and 2 µL of 10 × DNase I reaction buffer. Samples were incubated at 37 °C for 30 min, followed by capsid disruption and DNase inactivation at 95 °C for 10 min, then held at 4 °C. DNase-treated samples were serially diluted 10-fold in DNA suspension buffer (10 mM Tris, pH 8.0, 0.1 mM EDTA (T0223, Teknova Inc., Waukegan, IL, USA), 100 µg/mL poly[A] (10108626001, Sigma-Aldrich Corporation, St. Louis, MO, USA) and 0.01% Pluronic F-68 (24040032, ThermoFisher) to achieve concentrations within the optimal detection range. qPCR reactions were performed using GoTaq qPCR mix (Promega, Madison, WI, USA) with the following cycling conditions: 95 °C 2 min, followed by 45 cycles of 95 °C for 15 s and 60 °C for 60 s and reported as genomes per milliliter (vg/mL).

### Impurity quantification

Total DNA concentrations were quantified using a FluorGreen HyperLight kit (333-K1603, APExBio, Houston, TX, USA) according to the manufacturer’s protocol with a Synergy H1 plate reader (BioTek, Winooski, VT, USA). For the quantification of overall proteins after purification, a bicinchoninic acid (BCA) protein assay, Pierce (23227, ThermoFisher) was used according to the manufacturer’s instruction.

### Anion exchange high-performance liquid chromatography (AEX-HPLC)

AEX-HPLC was carried out on an Agilent 1260 Infinity II Bio-inert LC System equipped with diode array and fluorescence detectors, as described earlier [25]. Filled capsid ratios were determined using a BioPro IEX QF column (YMC CO., LTD, Kyoto, Japan, 30 × 4.6mm, 5 µm) at 25 °C with a flow rate of 0.6 mL/min. Separation was achieved using a Tris/ MgCl₂ buffer system and a stepwise gradient from 5% to 100% buffer B over 15 min.

### Transduction assay

Transduction efficiency of rAAV5 were determined using a GFP-based transduction assay in adherent HEK293 cells (CRL-1573, ATCC, Manassas, VA, USA). Cells were seeded at 3 × 10^5^ cells/mL in 24-well plates and incubated for 24 h. Prior to infection, 3 wells were sacrificed to determine VCC and culture medium was removed. Vector samples were normalized to equal genomic titers (vg/mL) and 150 µL of vector dilution in DMEM supplemented with 10% heat-inactivated FCS was added at a multiplicity of infection (MOI) of 2 × 10^4^ vg/cell. Cells were incubated with virus for 4 h, after which 500 µL of fresh DMEM medium was added, and cells were cultured for 72 h. Following, cells were harvested, strained (431752, Corning Inc., Corning, NY, USA) and analyzed by flow cytometry (SH800ZFP, Sony, San Jose, CA, USA). The transduction activity was calculated by dividing the absolute number of GFP-positive cells (TU, Transducing Unit) by the total number of viral genomes loaded per well.

### Process performance evaluation

The upstream comparison between batch and integrated processes were evaluated with respect to yield, accumulated product concentration and space time yield adapted from [33]. These three metrics can be calculated with respect to capsid and viral genomes. For the total capsid titer related yield, *Y_i_*, the following formula is used:

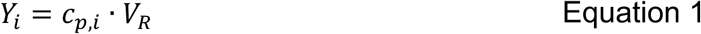

Where the *c_p_*_,*i*_ represents product concentration at time *i,* and *V_R_* corresponds to the working volume of the bioreactor. For the perfusion processes, the corresponding yield is calculated as follows:

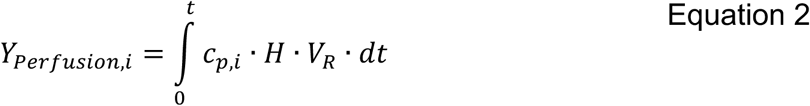

Where *H* denotes the perfusion rate. The corresponding space time yield *STY_i_* is calculated using 3:

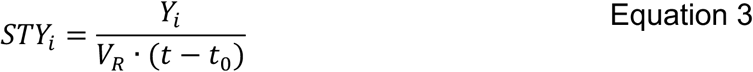

Sieving (*S*) was calculated after hollow fiber of the rTFF and sterile filtration as follows:

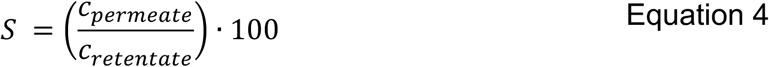

While *c_permeate_* represents the concentration of capsids particles after filtration and *c_retentate_* before filtration. The yield of the DSP capture, *Y_DSP_* was calculated using the following equation:

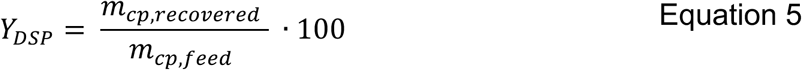

Here, *m_cp_*_,*recovered*_ represents the recovered product mass in capsid particles and *m_cp_*_,*feed*_ the total feed mass. The productivity, *P_DSP_* of the capture was determined as ratio of *m_cp_*_,*recovered*_ divided by the total time of the integrated downstream process, *t_totalDSP_* multiplied by the total column volume *V_column_*.

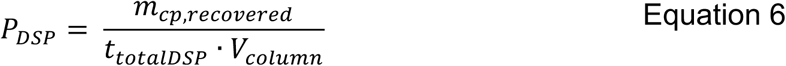

Buffer consumption, *BC* during the capture operation was calculated by dividing the total buffer volume, *V_buffer_* by the total recovered mass, *m_cp_*_,*recovered*_:

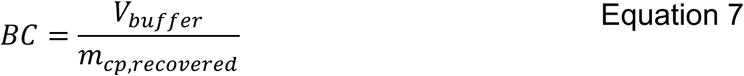

Performance indicators across batch and continuous processing modes were compared using an independent two-sample t-test, with the significance threshold set at p < 0.05.

## Results

### Experimental and process setup

The primary objective of this study was to demonstrate the feasibility of integrating a perfusion process with continuous multi-column capture for the production of rAAV. Whilst the majority of batch processes for rAAVs comprise a series of manual steps, and in our case include lysis, centrifugation, endonuclease treatment, sterile filtration and batch affinity chromatography, the proposed integrated continuous process requires fundamental process decisions to achieve continuous flow over an extended period (Figure 1).

Firstly, the decision was taken to capture exclusively extracellular rAAV, a strategy that reduces the total number of particles that can be captured. This sacrifice can be justified for most serotypes, given that the prolonged cultivation time in perfusion can significantly increase the total extracellular rAAV [16,17].

Secondly, the omission of the lysis step and endonuclease constitutes a further fundamental alteration to the process. As demonstrated in earlier studies on continuous multi-column capture, novel AAV affinity resins have been shown to be capable of effectively reducing DNA from non-lysed perfusion material [25].

Thirdly, it is imperative to acknowledge the significance of the sterility barrier and flow control, which are inextricably linked to the filtration strategy. Based on prior knowledge from antibody production, a scalable hollow fiber filtration technology for preliminary filtration and clarification using 0.4 µm in reverse TFF was implemented [14,34,35]. This was followed by two additional sterile filtrations before and after the surge tank. In order to maintain stability and sterility for all integrated runs, fixed flow rates were selected, and the sampling of the surge tank was a simple and effective way to control small flow rate perturbations.

Despite the fundamental differences between batch and integrated continuous processes, an objective comparison between the two approaches was attempted with two distinct processes for rAAV5 production (Process A and B). The primary difference between them is the choice of helper plasmid system (JD171 for Process A or pADΔF6 for Process B) and the insert (pCMV-GFP, while pProA23-GFP was used for early runs of Process A, see supplemental information Figure S1) utilized for triple-plasmid transient transfection.

### Cell culture process performance

To evaluate the performance of Process A and Process B under different cultivation strategies, viable cell concentration (VCC) and cell viability were monitored over time. The data of the batch and integrated continuous processes used for direct comparison are shown in Figure 2, while the complete data set from previous upstream runs are provided in supplemental information Figure S1 and Figure S2. The batch mode resulted in limited cell growth, with VCC remaining below 3 × 10^6^ cells/mL and the cultures being harvested 3 days post-transfection in all processes. In contrast, the perfusion mode supported robust cell growth post-transfection, with VCC steadily increasing to peak values between (50 –60) × 10^6^ cells/mL on day 8 for both processes. Transfection at day 0 induced a temporary drop in cell viability. While batch viability remained low after transfection at around 60 –90% in Process A and 80% in Process B, the integrated runs demonstrated a strong recovery, with viability climbing back up to exceed 95% by the end of the culture period. The integrated operation also yielded distinct differences in capsid production and secretion dynamics between the two processes. In Process A, a relatively stable extracellular capsid concentration between (2 –4) × 10^11^ cp/mL throughout day 3 to 8 was achieved. Comparing the resulting cumulative harvested product per volume cell culture in Figure 2 G (filled blue circles) is higher compared to harvested product in batch process (grey triangle in Figure 2 E).

**Figure 2.**
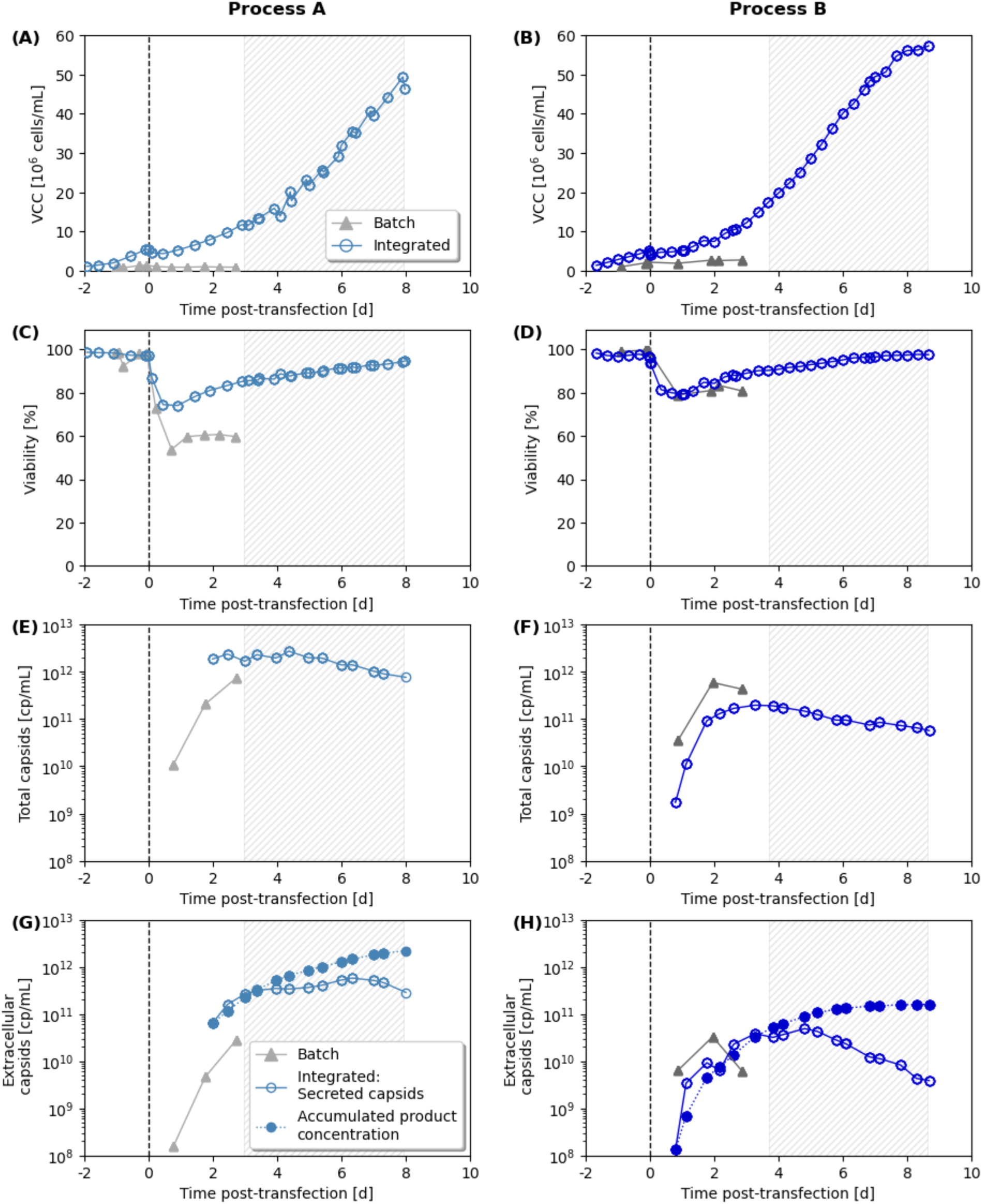
Comparison of upstream process performance between batch and perfusion for Process A (left panels) and Process B (right panels). Temporal profiles of viable cell concentration (A, B) and cell viability (C, D) demonstrate sustained cell growth and viability recovery post-transfection (vertical dashed line at t = 0) in the perfusion mode (blue circles) compared to the batch runs (grey triangles). Total capsid concentration (E, F) and extracellular capsid dynamics (G, H) highlight almost steady-state productivity during the integrated continuous harvest phase (shaded regions), showing both daily secreted capsids (open circles) and cumulative harvested product per volume cell culture (filled circles).

The integrated process B showed a lower total capsid concentration compared to Process A and also to a previously conducted perfusion Process B (see Figure S1), resulting in an extracellular capsid concentration of (0.5 –4) × 10^11^ cp/mL. Nevertheless, the mean extracellular accumulated harvested capsid per liter cell culture volume is higher in perfusion mode compared to the batch mode in both processes. On average, extracellular capsid in perfusion was for Process A and B: (1.90 ± 1.89) × 10^15^ compared to batch with (8.70 ± 4.25) × 10^14^.

Table 1 summarizes the upstream performance of batch and perfusion for Processes A and B based on capsid (cp) and vector genome (vg) production for all conducted cell culture processes. In Process A, perfusion significantly increased the total capsid yield by almost 4-fold (p<0.001) and resulted in a 58% (p=0.1) higher capsid space-time yield (STY) despite the prolonged cultivation time compared to batch processing. It is worth mentioning that the process time for the space-time yield includes the cell-growth-phase until reaching the required cell concentration for transfection and ends at the last harvest time point. In Process B, both modes yielded similar final capsid amounts; in contrast, perfusion achieved higher vector genome yields due to a higher full/empty ratio. Consequently, capsid-based STY was nearly twice as high for batch as for perfusion, whereas vector genome-based STY was practically identical between the two modes (see supplemental information Table S2 for results of individual processes). While Process A represents a case with high capsid titers, the proportion of ssDNA-filled particles measured by qPCR/ELISA was below 1% and was therefore deemed negligible. Consequently, the analysis of the CQAs was focused exclusively on Process B hereafter.

**Table 1.**
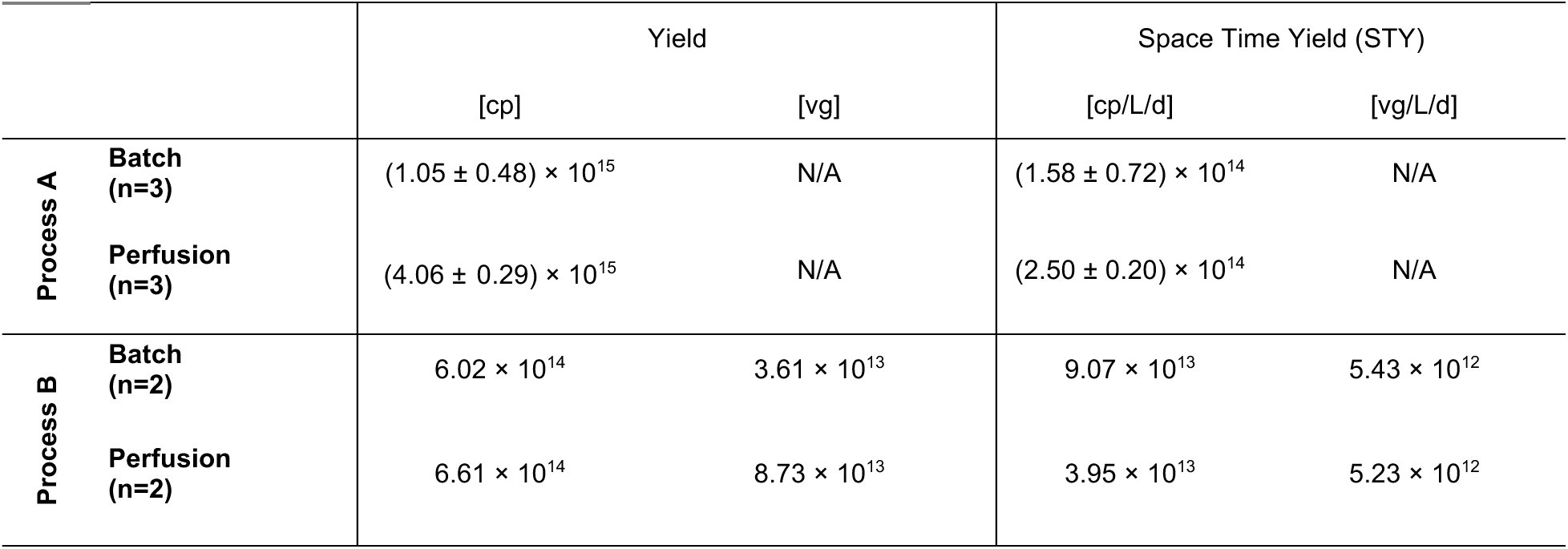
Comparison of upstream performance of batch and perfusion processes. Comparison of upstream performance between batch and integrated continuous operations for both process designs (A and B), with overall yield per bioreactor run (capsids and for vector genomes) and space time yield (STY) with respect to capsids per bioreactor volume and time and vector genomes per bioreactor volume and time.

|  |  | Yield |  | Space Time Yield (STY) |  |
| --- | --- | --- | --- | --- | --- |
|  |  | [cp] | [vg] | [cp/L/d] | [vg/L/d] |
| Process A | Batch (n=3) | $(1.05 \pm 0.48) \times 10^{15}$ | N/A | $(1.58 \pm 0.72) \times 10^{14}$ | N/A |
| | Perfusion (n=3) | $(4.06 \pm 0.29) \times 10^{15}$ | N/A | $(2.50 \pm 0.20) \times 10^{14}$ | N/A |
| Process B | Batch (n=2) | $6.02 \times 10^{14}$ | $3.61 \times 10^{13}$ | $9.07 \times 10^{13}$ | $5.43 \times 10^{12}$ |
| | Perfusion (n=2) | $6.61 \times 10^{14}$ | $8.73 \times 10^{13}$ | $3.95 \times 10^{13}$ | $5.23 \times 10^{12}$ |

### Filtration process performance

Given that the integrated perfusion-based rAAV production process is relying exclusively on extracellular particles, efficient continuous harvesting with minimal product loss is critical for maximizing overall yield. One potential limitation of prolonged hollow fiber membrane operation is the retention of rAAV particles, which could progressively reduce harvest efficiency over time. To assess this effect, capsid recovery was evaluated following primary clarification by rTFF hollow fiber filtration and subsequent sterile filtration across both integrated production processes A and B (Figure 3). A strong overlap between all three sampling positions was observed at each time point, indicating minimal product sieving during clarification (Figure 3 A, B). Mean capsid recovery after rTFF hollow fiber clarification reached 93 ± 8% and 92 ± 13% for Processes A and B, respectively, demonstrating negligible particle retention by the membrane (Figure 3 C and D). Subsequent 0.2 µm sterile filtration resulted in only minor additional losses, with final recoveries of 95 ± 11% for Process A and 86 ± 24% for Process B.

**Figure 3.**
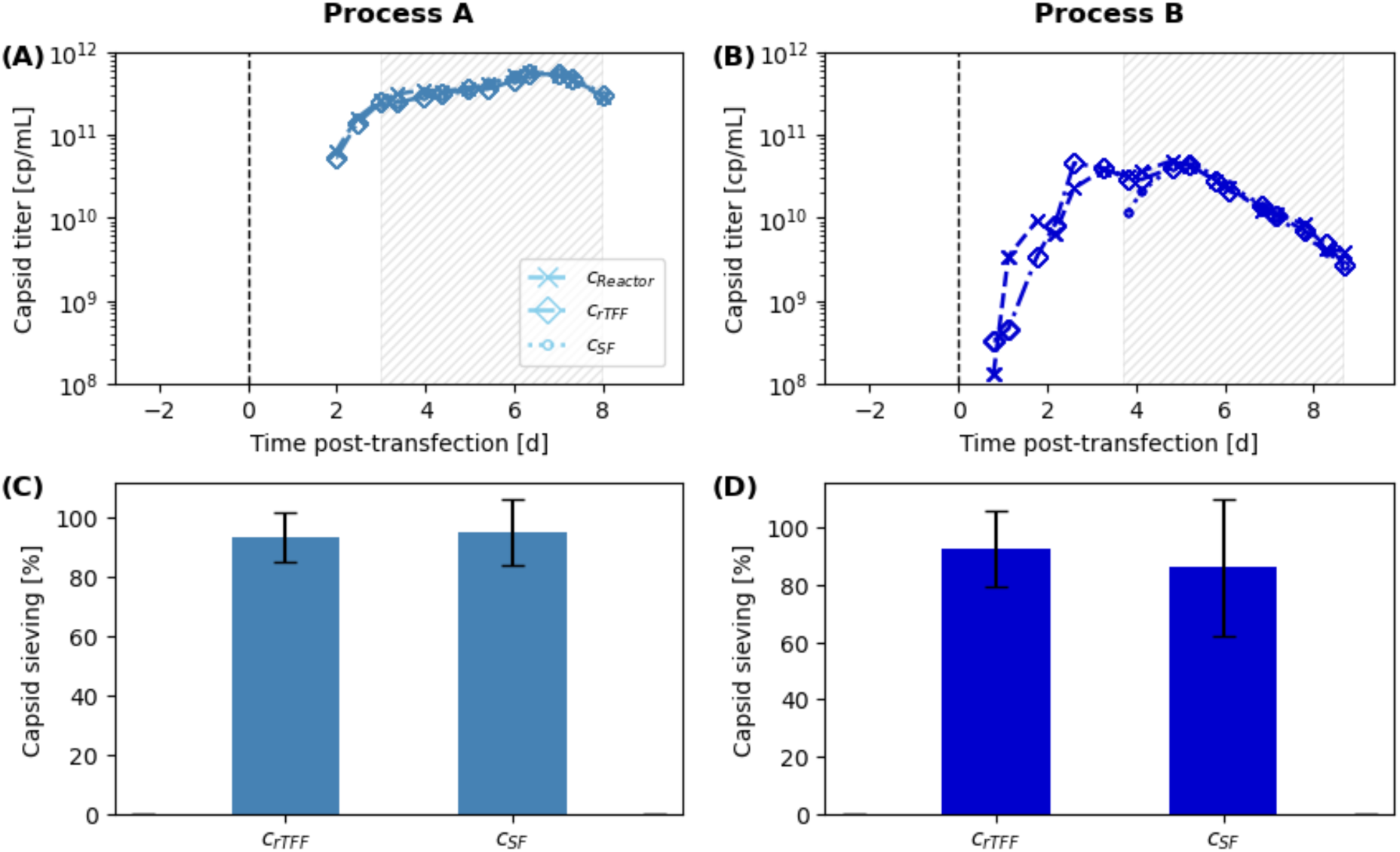
Filtration performance of rTFF and sterile filtration. Capsid concentration profiles are shown in the perfusion bioreactor (*c_Reactor_*), after the cell retention device (*c_rTFF_*) and after the 0.2 µm sterile filtration (*c_SF_*) step for the continuous integrated Process A (A) and process B (B), respectively. Average capsid sieving for rTFF and SF steps is shown in (C) for Process A and (D) for Process B. Standard deviations (error bars) were calculated across the entire process duration.

It is worth noting that a second sterile filtration was carried out after the surge tank to ensure sterility (Figure 1). No sampling was performed after this second sterile filtration. Therefore, a potential loss will be attributed to the continuous affinity chromatography step.

### Twin-column affinity purification

The variable titers over the multiple days of cultivation and filtration required adjustments to maintain a constant load without product loss in the flow-through during counter-current twin-column chromatography. This was achieved by limiting the total particles loaded during each of these phases to the same amount applied in the model CaptureSMB run as previously reported [25]. To calculate the interconnected loading time and the batch load flow rate, ELISA titers from previous non-integrated perfusion runs of the same process were used (Figure S1). The integrated capture of Process A included 17 cycles (Figure 4 A). The peaks showed comparable peak height and shape, as seen in the overlaid elution profiles (supplemental information Figure S4). This indicated that the load adjustment based on historical ELISA titers was adequate to maintain a constant load amount. Additionally, the high similarity suggests that there was no fouling over the course of the operation.

**Figure 4.**
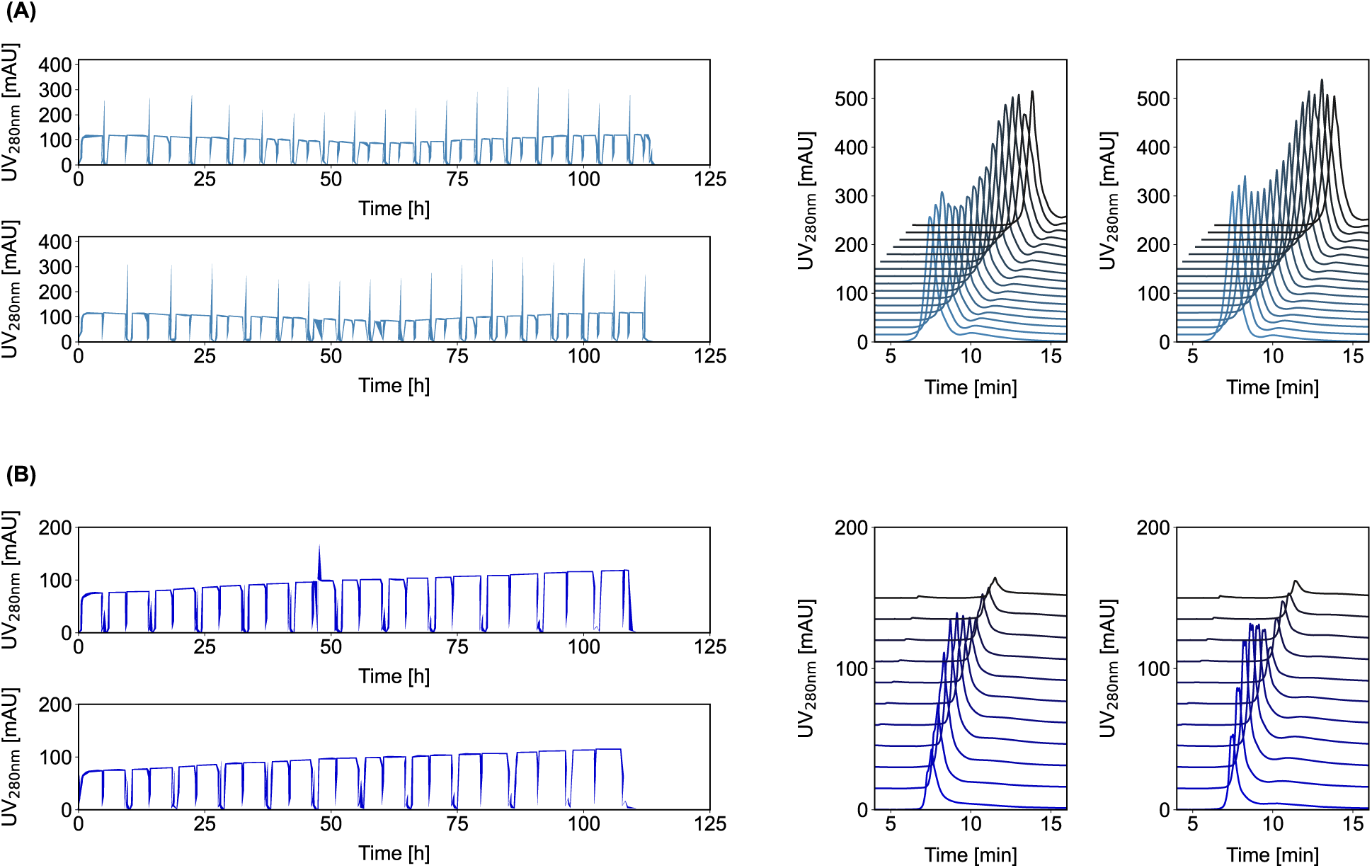
UV_280nm_ profiles for the two integrated continuous chromatography processes. **A and B**. (A) Process A (17 cycles) and (B) Process B (11 cycles) are shown. The left panels depict the continuous UV profiles for column 1 (top) and column 2 (bottom). The right panels display the overlaid elution peaks for column 1 (left) and column 2 (right), with progressively darker colors indicating later cycles.

The same adjustments were scheduled for the integrated capture of Process B. However, the capsid titers were much lower in comparison to Process A and also lower compared to the previously non-integrated perfusion Process B run. Based on previous results from the rAAV CaptureSMB operations, extended loading phases may result in faster column fouling [25]. Consequently, the maximal interconnected load time was limited rather than the maximal particle load to prevent premature fouling. To ensure the interconnected operation can be maintained for a comparable duration as in Process A, the load duration was limited to 3.3 h and was only extended to 4.3 h for the last three cycles, where the titers had already declined substantially. This adjustment led to substantially smaller elution peaks and negatively impacted process performance (Figure 4 B and Figure S3 B). Nonetheless, the strategy ensured a successful integrated capture for 11 cycles (22 switches) for up to five days without signs of column fouling.

The performance of the continuous DSP capture was further assessed by comparing the yield, buffer consumption and productivity of the continuous process with those of a conventional batch capture. For this purpose, the integrated processes were compared with conventional batch capture runs, using lysed stirred-tank reactor feed material. Yield, buffer consumption and productivity were calculated for total particles, and in the case of Process B, additionally for full particles. Process A represented a scenario with high capsid titers, where adjustment of the interconnected and batch load to the target particle load was possible. In this comparison, the yield (- 24%) of the continuous process did not improve compared to the conventional batch. In contrast, the buffer consumption per recovered particle was reduced by - 71%, and the productivity slightly increased by + 5% (Table 2 and supplemental information Table S3).

**Table 2.** Comparison of downstream performance of batch and integrated continuous processes. Comparison of downstream performance between batch and integrated continuous operations for both process designs (A and B), with respect to load density, yield, productivity, and buffer consumption. For integrated operations, all calculations include the start-up and shutdown phases.

|  |  | Number of switches | Column volume | Load density | Yield |  | Productivity |  | Buffer consumption |  |
| --- | --- | --- | --- | --- | --- | --- | --- | --- | --- | --- |
|  |  | [#] | [mL] | [cp/mL resin] | [% cp] | [% vg] | [cp/L/h] | [vg/L/h] | [L/cp] | [L/vg] |
| **Process A** | **Batch (n=2)** | 1 | 0.491 | $2.21 \times 10^{16}$ | 89.9 | N/A | $9.81 \times 10^{15}$ | N/A | $4.02 \times 10^{-15}$ | N/A |
| | **Integrated (n=2)** | 28 / 32 | $2 \times 0.491$ | $5.47 \times 10^{16}$ | 68.0 | N/A | $1.03 \times 10^{16}$ | N/A | $1.16 \times 10^{-15}$ | N/A |
| **Process B** | **Batch (n=2)** | 1 | 0.491 | $2.36 \times 10^{16}$ | 83.9 | 75.4 | $9.56 \times 10^{15}$ | $4.63 \times 10^{14}$ | $4.04 \times 10^{-15}$ | $8.48 \times 10^{-14}$ |
| | **Integrated (n=1)** | 22 | $2 \times 0.491$ | $6.99 \times 10^{15}$ | 71.3 | 82.3 | $9.94 \times 10^{14}$ | $1.32 \times 10^{14}$ | $1.41 \times 10^{-14}$ | $1.07 \times 10^{-13}$ |

By comparison, the capsid titers for Process B were considerably lower, although a higher full/empty ratio was achieved. Consequently, the columns were not fully loaded to the target total particle amount. This was reflected in the much lower productivity (- 90%) and higher buffer consumption (+ 250%), as well as the lower yield (-15%) compared to conventional batch chromatography (Table 2). Due to the low capsid titers and the risk of premature column fouling, the capsid load was reduced to maintain process operation, which negatively impacted the performance. When the same process comparison was carried out for the full AAVs only, the productivity (-71%) and buffer consumption (+26%) remained inferior to the conventional batch process, yet the vector genome yield improved, showing a +9% increase (Table 2).

### Product recovery and DNA reduction

The performance of the integrated Process B against the traditional batch process is shown in Figure 5, where product recovery and total DNA reduction were monitored across key unit operations. Both the batch (grey) and integrated continuous (blue) processes maintained stable, near-complete product recovery through the early stages of purification. As the processes progressed, cumulative product recovery gradually decreased to similar final levels. Interestingly, the measured titers increased after endonuclease treatment for the batch processes and thus the titers in the batch process might have been slightly underestimated. Potential reasons for that increase before fragmentation of host-cell DNA might be effects caused by aggregation of DNA or competition for polymerase binding [36]. Following the chromatography step, the batch process yielded a product recovery of approximately 67%, while the integrated process achieved a comparable recovery of approximately 60%.

**Figure 5.**
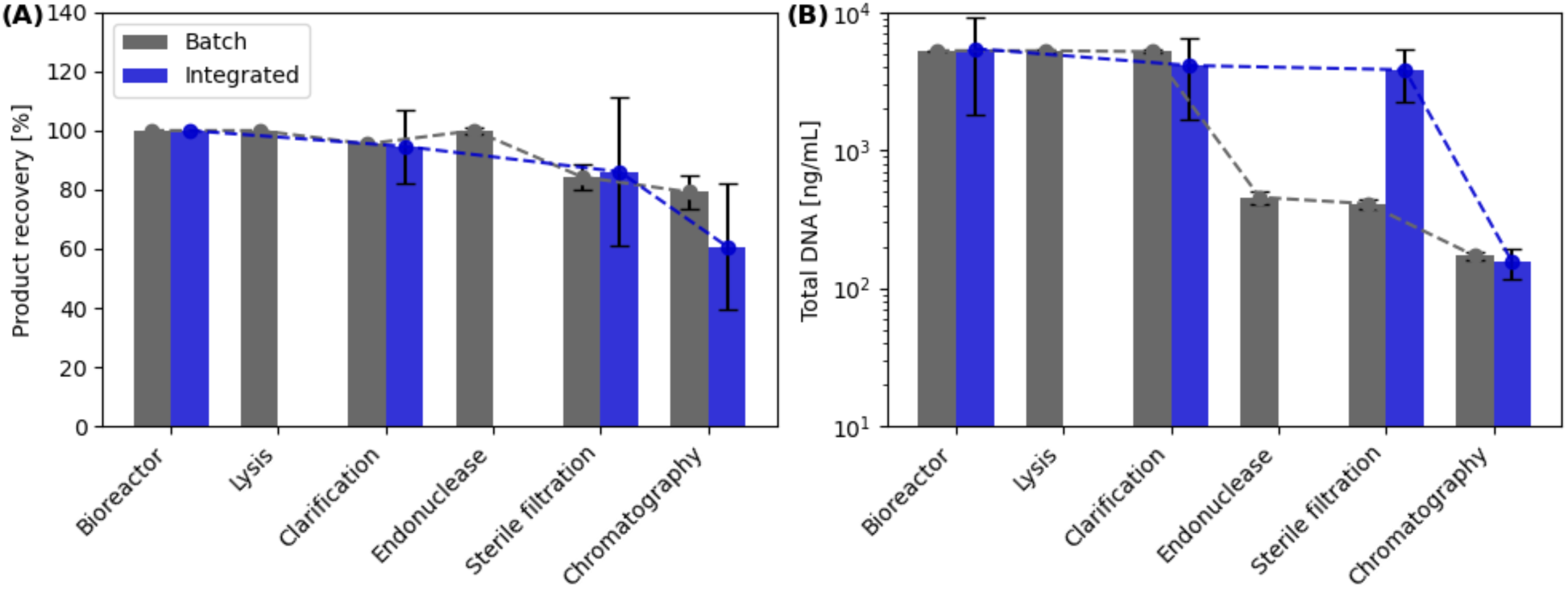
Comparison of product recovery and DNA reduction between batch and integrated continuous Process B. (A) Product recovery (%) and (B) total DNA concentration (ng/mL) compared across the unit operations of the batch (grey) and integrated (blue) processes. Unit operations for the batch process include bioreactor, lysis, clarification, endonuclease treatment, sterile filtration, and chromatography, whereas the integrated continuous process excludes the lysis and endonuclease stages. For the batch process, error bars denote the standard deviation of duplicate runs; for the integrated process, they reflect the standard deviation of yields evaluated across the process. The value of the endonuclease product recovery of the batch exceeded the initial bioreactor concentration and was corrected to 100%.

The total DNA concentration profiles highlight the structural differences between the two processes. In the batch process, DNA levels peaked following the lysis step (reaching nearly 10^4^ ng/mL) yet were dramatically reduced by more than one order of magnitude during the endonuclease treatment (decreasing to approximately 5 × 10^2^ ng/mL). In contrast, the integrated continuous process, which bypasses both the discrete lysis and endonuclease stages, maintained a relatively high and steady DNA concentration (around 5 × 10^3^ ng/mL) through the sterile filtration step. The high initial DNA content during the integrated continuous process can be explained by the high cell concentration in the perfusion bioreactor. Remarkably, the chromatography step in the integrated continuous process proved highly effective, reducing the total DNA concentration down to approximately 10^2^ ng/mL. This final concentration is comparable to, or even slightly lower than, the final DNA level achieved by the batch process, thereby demonstrating that the twin-column chromatography can successfully resolve the DNA burden even without upstream endonuclease treatment.

### Process and product quality consistency

To investigate the impact of prolonged integrated operation on product quality, critical quality attributes (CQAs) of the eluted rAAV5 particles from Process B were analyzed across the capture switches and compared with the batch purified samples (Figure 6). The step recovery as well as total protein content remained generally stable throughout the entire integration period, while total DNA showed a slight increase over time but remained, on average, below the value observed for the batch process. Furthermore, AEX-HPLC showed a consistent filled capsid ratio of 29.1 ± 1.2% for the integrated process compared to 26.7 ± 0.3% for the batch capture (Figure 6 D). Moreover, the full/empty capsid ratio was determined using qPCR/ELISA, resulting in a mean ratio of 9.19 ± 4.41% for the integrated process compared to 6.8 ± 0.2% for the batch process (Figure 6 E). Compared with AEX-HPLC, the qPCR/ELISA-based method showed substantially higher variability, and a decline of filled particles after switch 12. To assess the functional quality of the eluted rAAV5 vectors, potency was compared between the integrated Process B and the corresponding batch (Figure 6 F). Across the 22 switches, the integrated process showed a mean specific activity of (5.1 ± 1.6) × 10^-5^ TU/vg, compared to (4.2 ± 1.6) × 10^-6^ TU/vg for the batch process. For visualization purposes, relative potency values are shown, with specific activity being normalized to the mean of batch specific activity.

**Figure 6.**
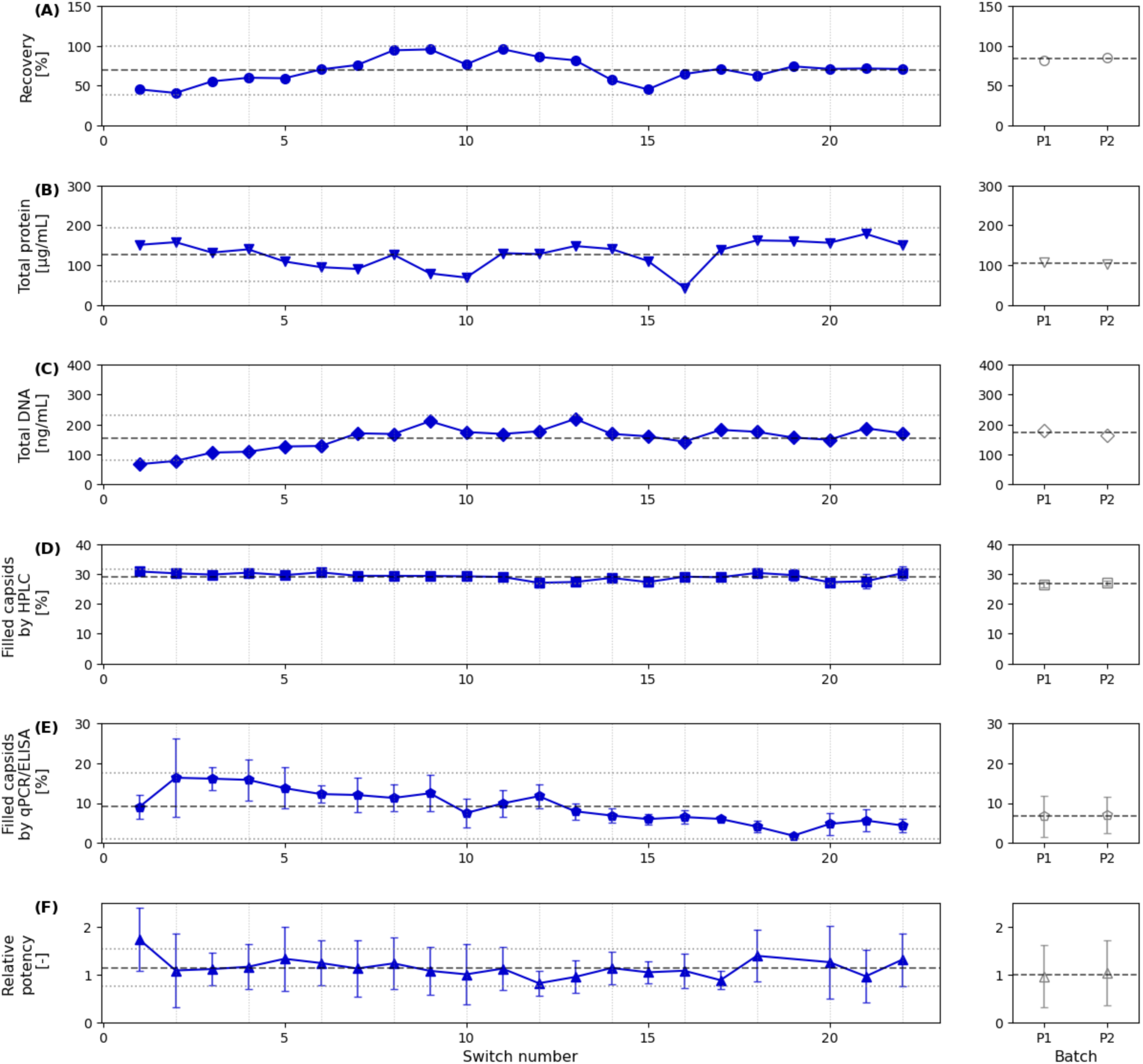
Comparison of performance indicators and product quality attributes of Process B batch and integration. Continuous process profiles across 22 column switches (left, blue) are compared against duplicate batch runs, P1 and P2 (right, grey, with dashed and dotted lines indicating batch mean with analytical standard deviation). Process performance is evaluated by (A) product recovery, (B) total protein, and (C) total DNA. Product quality is assessed via filled capsid percentages determined by (D) HPLC and (E) qPCR/ELISA, alongside (F) relative potency. Results from relative potency of switch number 19 were excluded as an outlier with a relative potency > 2. Error bars represent the standard deviation of analytical triplicate measurements.

## Discussion

This study demonstrates the successful integration of upstream perfusion cultivation and continuous two-column affinity capture into a stable continuous biomanufacturing workflow for rAAV production. Beyond the technical integration itself, one of the central achievements of this work was the ability to maintain stable and sterile operation over several days of uninterrupted processing [37]. Maintaining sterility is a critical requirement for any integrated continuous biomanufacturing process, particularly when multiple unit operations and filtration steps are directly connected and operated under low flow conditions over extended time periods [38]. The combination of rTFF clarification with sequential 0.2 µm sterile filtration and a controlled surge tank strategy proved sufficient to establish a robust sterility barrier throughout the integrated process. No operational instability or indications of contamination were observed during any of the integrated runs, highlighting the practicality of this architecture for longer-term operation.

A major conceptual shift of the presented workflow compared to conventional rAAV manufacturing lies in the omission of several standard upstream and harvest operations. Traditional batch manufacturing typically relies on cell lysis, clarification by centrifugation and/or depth filtration, endonuclease treatment, and many batch processes include intermediate concentration steps prior to affinity purification [39]. With cell lysis being the step introducing high burden on downstream purification by releasing a substantial amount of host cell DNA and HCPs [40]. In contrast, the present approach directly harvests extracellular rAAV particles from perfusion culture and transfers them continuously to downstream capture. Furthermore, the use of extracellular material has been reported to be advantageous over lysed material for subsequent affinity purification [41]. By avoiding lysis, the release of intracellular DNA and host cell impurities is substantially reduced, thereby simplifying downstream processing. The elimination of endonuclease treatment did not compromise final DNA reduction, as the affinity capture step alone was sufficient to reduce DNA concentrations to levels comparable to the conventional batch process. This finding is particularly relevant given the significant material costs associated with nuclease treatment in current rAAV workflows [23]. In fact, endonuclease-free clarification processes may result in >90 % reduction in cost-of-goods of the upstream harvesting step [24].

The omission of centrifugation and depth filtration further streamlines the process and may reduce product losses commonly associated with clarification operations. Previous studies have reported substantial rAAV losses during clarification, especially for fragile viral vectors and at larger process scales [23]. In the present work, the rTFF-based clarification strategy achieved consistently high capsid recoveries above 90% over multiple days of operation. These results demonstrate that hollow-fiber based clarification can provide an effective alternative to conventional clarification technologies for integrated continuous viral vector manufacturing. For production of smaller biologics such as monoclonal antibodies, high product-sieving cell retention systems were reported and are essential for extended runtime and efficient harvest [34,42]. Moreover, no significant decline in sieving performance was observed over time, indicating that membrane fouling and product retention remained limited despite prolonged exposure to high-cell-density cultures and cell-derived impurities.

The integrated twin-column affinity capture process demonstrated stable operation throughout the cultivation period, indicating that CaptureSMB can be directly coupled to continuously harvested perfusion material. While previous studies established the applicability of CaptureSMB for rAAV purification under non-integrated conditions [25], the direct integration of continuous affinity chromatography with fluctuating perfusion harvest streams remained largely unexplored. Despite continuous feeding over up to five days, no progressive loss in column performance, breakthrough control, or elution reproducibility was observed, indicating that the system tolerated prolonged exposure to crude harvest material without signs of fouling. In Process A, high upstream particle titers enabled operation near the target column loading and produced highly consistent elution profiles across all switches, demonstrating that robust synchronization between fluctuating USP production and DSP capture is achievable. These findings support that two-column affinity capture represents a viable strategy for fully integrated continuous rAAV DSP.

At the same time, the Process B experiments highlighted one of the key operational challenges for integrated rAAV manufacturing: the dynamic nature of viral vector titers during prolonged cultivation. In contrast to monoclonal antibody perfusion processes, with reported stable steady-state productivity during prolonged operation [34], transiently transfected rAAV production systems exhibit stronger temporal fluctuations in extracellular particle concentrations due to transient transfection dynamics, shifts in viability, changing secretion behavior as well as heterogeneity [43]. Consequently, maintaining consistent column loading conditions becomes considerably more challenging. In this study, historical titer profiles were used to estimate interconnected load durations and batch load flow rates. While this strategy enabled straightforward process integration, deviations between expected and actual titers negatively affected column utilization and process productivity, effects that were particularly evident in the integrated continuous Process B, highlighting a common challenge associated with extended integrated continuous bioprocess operation. In the present study, Process B exhibited a substantially stronger titer variation, with a maximum titer increase of approximately 162% over the course of integrated operation, further highlighting the operational challenges associated with upstream titer variability and its direct impact on downstream performance during prolonged integrated continuous processing.

These observations emphasize the necessity for real-time or near real-time analytical monitoring strategies for future industrial implementation. In particular, process analytical technology (PAT) solutions capable of rapidly quantifying rAAV titers would substantially improve process control and dynamic loading adjustments during continuous capture [44]. Online HPLC-based methods or UV-based dynamic control could enable adaptive control of loading durations, flow rates, and switching intervals based on actual process conditions rather than historical estimates [45,46]. Such approaches would improve column utilization, minimize underloading or breakthrough risks, and increase overall process robustness. In addition, integration of PAT frameworks with supervisory process control systems could facilitate automated operation and support the transition toward fully autonomous continuous manufacturing platforms.

While this study focused on rAAV5 production using transient transfection, the modular architecture of this integrated process is highly amenable to expansion. The transient transfection-based process has its bottleneck by the time span of rAAV expression as well as level of secretion depending on the serotype and construct [47]. To favor the secretion pathway of rAAV, optimization of expression and sequence of the membrane-associated accessory protein (MAAP) could be considered also to utilize low-secreting serotypes such as rAAV2 [48,49]. Transitioning from transient transfection to stable or inducible producer cell-lines may result in more consistent titer and potentially even eliminate the temporal titer fluctuations. Assuming that upstream yields continue to exponentially increase as has been the case in the last two decades, intensification of downstream unit operations will become increasingly important. One limitation of the present study is that the continuous downstream process was somewhat oversized relative to the achieved titers; higher upstream titers would be expected to further improve overall process performance of the integrated continuous compared to the batch process.

It is also worth noting that an integrated continuous approach may never truly achieve a steady state, or even a quasi-steady state. As observed in certain antibody production processes, product concentration can vary substantially, and dynamic control of column loading can adequately accommodate such fluctuations. Indeed, non-steady-state integrated continuous processes, in which all product fractions are pooled, may arguably represent a more pragmatic approach, as batch definition, final product quality assessment, and consequently regulatory filing are considerably more straightforward.

Furthermore, this integrated perfusion-to-capture configuration holds substantial potential for highly unstable viral vectors, such as envelope-pseudotyped lentiviruses. These vectors are notoriously sensitive to environmental parameters and enzymatic degradation, exhibiting half-lives of only a few hours under physiological conditions [9]. By continuously harvesting lentiviruses or other fragile viral vectors via perfusion and immediately capturing them on a twin-column system, the overall residence time of the fragile vector within the crude bioreactor environment is drastically reduced.

## Concluding remarks

This study establishes a proof-of-concept for an integrated, continuous biomanufacturing platform for rAAV production. By coupling rTFF-based perfusion with a twin-column CaptureSMB system, this process bypasses several traditional, yield-limiting, and cost-intensive batch steps, including cell lysis, depth filtration, endonuclease treatment, and intermediate concentration, while maintaining final product quality, recovery, and purity over several days of uninterrupted operation. Although the dynamic titers associated with transient transfection introduce operational challenges that need to be addressed in future work, the overall design of the platform closely resembles established continuous antibody manufacturing schemes, supporting its potential scalability and industrial applicability.

Ultimately, this platform is well-positioned to leverage ongoing advances in improved rAAV secretion and inducible expression systems. Integrating these technologies has the potential to further enhance productivity and reduce buffer consumption, while providing a more stable and continuous viral vector supply to fully utilize the efficient purification capabilities of this process. In combination with advanced process monitoring and PAT-enabled control strategies, integrated continuous rAAV biomanufacturing may pave the way to substantially reduce production costs, improve process robustness, and enhance product consistency. Collectively, these advances establish continuous manufacturing as a scalable, economically sustainable framework critical to expanding access to life-saving gene therapies.

### Outstanding Questions

To what extent can the drastically shortened product residence times inherent to continuous biomanufacturing prevent deleterious capsid post-translational modifications, such as surface deamidation or oxidation, that typically accumulate during the prolonged incubation and harsh conditions of batch harvesting?

Can host cell engineering strategies - specifically the transition to stable, inducible producer cell lines - be synergistically combined with continuous harvesting to eliminate transient transfection variability and actively force extracellular secretion of poorly egressing serotypes?

How can non-destructive, real-time process analytical technology (PAT) sensors for online capsid quantification deployed to enable fully autonomous, closed-loop automated control of continuous viral vector processes?

What is the theoretical and practical maximum ceiling of this platform’s volumetric productivity and economic benefits?

Given that many gene therapies target ultra-rare orphan indications, is a decentralized “scale-out” manufacturing model of parallelized, small-scale continuous systems economically and regulatorily superior to traditional vertical “scale-up” in large-scale batch bioreactors?

## Acknowledgements

This study was supported by ChromaCon AG and YMC CO., LTD. We would like to thank the team of Levitronix Technologies, explicitly Philip Giller, Vito Buffa, Petrit Loshi and Patrick Romann for their suggestions, discussions and support for the rTFF setup. We would also like to thank the team of Asahi Kasei Bioprocesses, explicitly Konstantin Doshishti-Agolli, Andrey Katalevsky and Achraf Jazi for their support and helpful discussions. Further, we would like to thank Nova Biomedical for their support, explicitly Evgueni Voronkov and Royston Bulman.

## Author contributions

**R.P.**: Conceptualization and design, methodology, validation, formal analysis, investigation, data curation, writing (original draft), writing (editing and review), visualization. **J.M.M.**: Conceptualization and design, methodology, validation, formal analysis, investigation, data curation, writing (original draft), writing (editing and review), visualization. **D.T.**: Methodology, data curation. **E.S.**: Methodology, data curation. **P.W.**: methodology. **Y.H.**: writing (editing and review). **R.T.**: writing (editing and review). **S.V.**: writing (editing and review). **T.M.S.**: writing (editing and review). **S.G.**: Conceptualization and design, validation, formal analysis, data curation, writing (original draft), writing (editing and review), visualization. **T.K.V.**: Conceptualization and design, investigation, writing (original draft), writing (editing and review).

## Data Availability

Data is available upon reasonable request from the corresponding author.

## References

1. A Reid, C., et al. (2024) Advancing AAV production with high-throughput screening and transcriptomics. Cell Gene Ther. Insights 10, 821–840

2. Zwi-Dantsis, L. et al. (2025) Adeno-Associated Virus Vectors: Principles, Practices, and Prospects in Gene Therapy. Viruses 17, 239

3. Maguire, A.M. et al. (2019) Efficacy, Safety, and Durability of Voretigene Neparvovec-rzyl in RPE65 Mutation–Associated Inherited Retinal Dystrophy. Ophthalmology 126, 1273–1285

4. Malm, M. et al. (2020) Evolution from adherent to suspension: systems biology of HEK293 cell line development. Sci. Rep. 10, 18996

5. Fu, Q. et al. (2023) Critical challenges and advances in recombinant adeno-associated virus (rAAV) biomanufacturing. Biotechnol. Bioeng. 120, 2601–2621

6. Sarkis, M. et al. (2023) Characterization of key manufacturing uncertainties in next generation therapeutics and vaccines across scales. J. Adv. Manuf. Process. 5, e10158

7. Karst, D.J. et al. (2018) Continuous integrated manufacturing of therapeutic proteins. Curr. Opin. Biotechnol. 53, 76–84

8. Gupta, P. et al. (2021) Economic assessment of continuous processing for manufacturing of biotherapeutics. Biotechnol. Prog. 37

9. Klimpel, M. et al. (2023) Development of a perfusion process for continuous lentivirus production using stable suspension producer cell lines. Biotechnol. Bioeng. 120, 2622–2638

10. Solis Olivares, A., et al. (2026) Challenges and opportunities in continuous bioprocessing of lentiviral vectors and adeno-associated viral vectors. Biotechnol. Adv. 90, 108923

11. Bielser, J.-M. et al. (2018) Perfusion mammalian cell culture for recombinant protein manufacturing –A critical review. Biotechnol. Adv. 36, 1328–1340

12. Karst, D.J. et al. (2016) Characterization and comparison of ATF and TFF in stirred bioreactors for continuous mammalian cell culture processes. Biochem. Eng. J. 110, 17–26

13. Coffman, J. et al. (2021) A common framework for integrated and continuous biomanufacturing. Biotechnol. Bioeng. 118, 1735–1749

14. Romann, P. et al. (2024) Co-current filtrate flow in TFF perfusion processes: Decoupling transmembrane pressure from crossflow to improve product sieving. Biotechnol. Bioeng. 121, 640–654

15. Dhingra, A. et al. (2026) Mechanistic Modeling of Hollow Fiber Fouling and Sieving Predictions for Continuous Bioprocessing. Biotechnol. Bioeng. 123, 1364–1379

16. Mendes, J.P. et al. (2022) AAV process intensification by perfusion bioreaction and integrated clarification. Front. Bioeng. Biotechnol. 10, 1020174

17. Zhang, Y. et al. (2025) Intensification of rAAV Production Based on HEK293 Cell Transient Transfection. Biotechnol. J. 20

18. Gränicher, G. et al. (2021) A high cell density perfusion process for Modified Vaccinia virus Ankara production: Process integration with inline DNA digestion and cost analysis. Biotechnol. Bioeng. 118, 4720–4734

19. Benskey, M.J. et al. (2016) Continuous Collection of Adeno-Associated Virus from Producer Cell Medium Significantly Increases Total Viral Yield. Hum. Gene Ther. Methods 27, 32–45

20. Park, D., et al. (2024) Continuous Production of Recombinant Adeno-associated Viral Vectors via Transient Transfection of HEK293 Cells in Perfusion Bioreactor. In Computer Aided Chemical Engineering, pp. 2587–2592, Elsevier

21. Chahal, P.S. et al. (2014) Production of adeno-associated virus (AAV) serotypes by transient transfection of HEK293 cell suspension cultures for gene delivery. J. Virol. Methods 196, 163–173

22. Leibiger, T.M. et al. (2025) Characterization of difficult-to-remove host cell proteins in adeno-associated virus downstream processing. Mol. Ther. Methods Clin. Dev. 33, 101623

23. Yang, O. et al. (2023) Process Design and Comparison for Batch and Continuous Manufacturing of Recombinant Adeno-Associated Virus. J. Pharm. Innov. 18, 275–286

24. Thakur, G. et al. (2025) Single-use chromatographic clarification to eliminate endonuclease treatment in production of recombinant adeno-associated viral vectors. Sep. Purif. Technol. 354, 128557

25. Müller, J.M. et al. (2026) Continuous Capture of recombinant AAV Particles Using Twin-Column CaptureSMB. bioRxiv [Preprint]. 2026. Available from: 10.64898/2026.06.12.731701

26. Mahajan, E. et al. (2012) Improving affinity chromatography resin efficiency using semi-continuous chromatography. J. Chromatogr. A 1227, 154–162

27. Mendes, J.P. et al. (2022) Continuous Affinity Purification of Adeno-Associated Virus Using Periodic Counter-Current Chromatography. Pharmaceutics 14, 1346

28. Müller, J.M. et al. (2025) Enrichment of Full AAV2 Using Multicolumn Countercurrent Solvent Gradient Purification (MCSGP). Biotechnol. Bioeng. 122, 2420–2432

29. Ossi, L. et al. (2025) Separation of empty and full adeno-associated viral capsids by MCSGP: From model-based design to implementation. J. Chromatogr. A 1758, 466219

30. Jüttner, J. et al. (2019) Targeting neuronal and glial cell types with synthetic promoter AAVs in mice, non-human primates and humans. Nat. Neurosci. 22, 1345–1356

31. Werder, P. et al. (2026) Intensified Plasmid Capture Process via Continuous Lysis and Membrane Chromatography. SSRN [Preprint]. 2026. Available from: 10.2139/ssrn.7316549

32. Su, W. et al. (2022) Self-attenuating adenovirus enables production of recombinant adeno-associated virus for high manufacturing yield without contamination. Nat. Commun. 13, 1182

33. Bausch, M. et al. (2019) Recommendations for Comparison of Productivity Between Fed-Batch and Perfusion Processes. Biotechnol. J. 14, 1700721

34. Karst, D.J. et al. (2017) Process performance and product quality in an integrated continuous antibody production process. Biotechnol. Bioeng. 114, 298–307

35. Romann, P. et al. (2026) A General Workflow for Tangential Flow Filtration Perfusion Scale-Up. Biotechnol. Bioeng. [Preprint]. 2026. Available from: 10.1002/bit.70335

36. Long, S. (2022) In pursuit of sensitivity: Lessons learned from viral nucleic acid detection and quantification on the Raindance ddPCR platform. Methods 201, 82–95

37. Konstantinov, K.B. and Cooney, C.L. (2015) White Paper on Continuous Bioprocessing May 20–21 2014 Continuous Manufacturing Symposium. J. Pharm. Sci. 104, 813–820

38. Ito, T. et al. (2024) Implementation of filtration models for filter performance prediction in bioprocess dead-end membrane filtration. Biochem. Eng. J. 208, 109358

39. Jungbauer, A. and Wheelwright, S. (2025) Downstream processing of AAV based gene therapy vectors. Sep. Purif. Technol. 368, 133051

40. Dobrowsky, T. et al. (2021) AAV manufacturing for clinical use: Insights on current challenges from the upstream process perspective. Curr. Opin. Biomed. Eng. 20, 100353

41. Leibiger, T.M. et al. (2026) Harvest Process and Affinity Resin Selection Impacts on Adeno-Associated Virus Residual Host Cell Protein Retention. Hum. Gene Ther. 37, 600–608

42. Feidl, F. et al. (2020) Process-wide control and automation of an integrated continuous manufacturing platform for antibodies. Biotechnol. Bioeng. 117, 1367–1380

43. Ladd, B. et al. (2025) Heterogeneity in an adeno-associated virus transfection-based production process limits the production efficiency. Sci. Rep. 15, 38459

44. Gerzon, G. et al. (2022) Process Analytical Technologies –Advances in bioprocess integration and future perspectives. J. Pharm. Biomed. Anal. 207, 114379

45. Heckel, J. et al. (2025) Rapid At-Line AAVX Affinity HPLC: Enabling Process Analytical Technology for Bioprocess Development of Adeno-Associated Virus Vectors. Biotechnol. J. 20, e202400656

46. Fioretti, I. et al. (2024) UV-based dynamic control improves the robustness of multicolumn countercurrent solvent gradient purification of oligonucleotides. Biotechnol. J. 19, 2400170

47. Piras, B.A. et al. (2016) Distribution of AAV8 particles in cell lysates and culture media changes with time and is dependent on the recombinant vector. Mol. Ther. - Methods Clin. Dev. 3, 16015

48. Schieferecke, A.J. et al. (2024) Evolving membrane-associated accessory protein variants for improved adeno-associated virus production. Mol. Ther. 32, 340–351

49. Elmore, Z.C. et al. (2021) The membrane associated accessory protein is an adeno-associated viral egress factor. Nat. Commun. 12, 6239

