## Supplemental Information: for "Integrated Continuous Biomanufacturing of Recombinant Adeno-Associated Virus"

### Supplemental Information: Integrated Continuous Biomanufacturing of Recombinant Adeno-Associated Virus

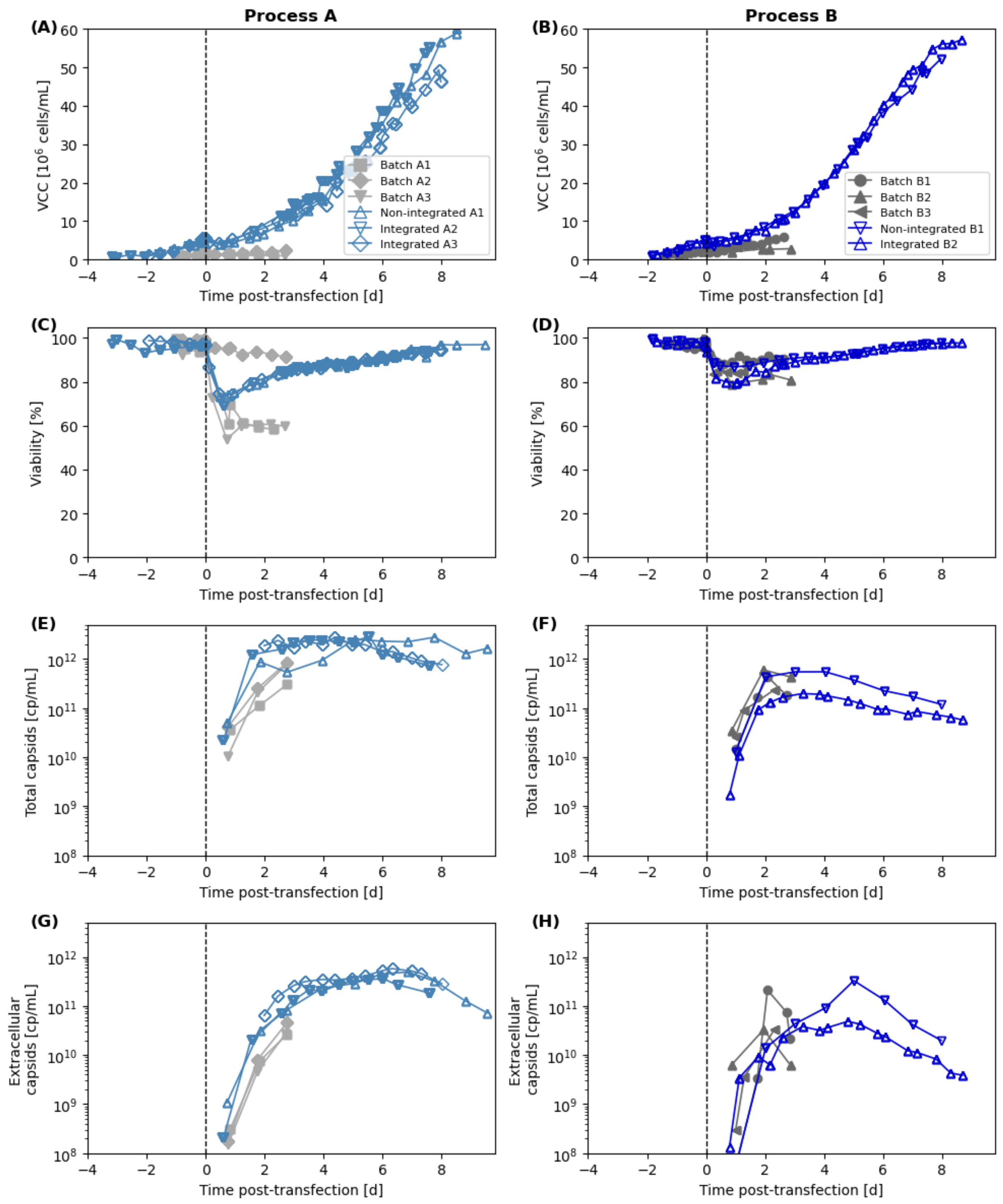


**Figure S1 Comparison of batch (grey-scale) and continuous process (integrated and non-integrated) (blue-scales) of Process A and Process B.** Time of transfection is indicated at time 0, with a vertical dotted line. A and B shows the viability of the processes, C and D visualizes the viable cell concentration. E and F the total capsids, G and H the secreted capsids.

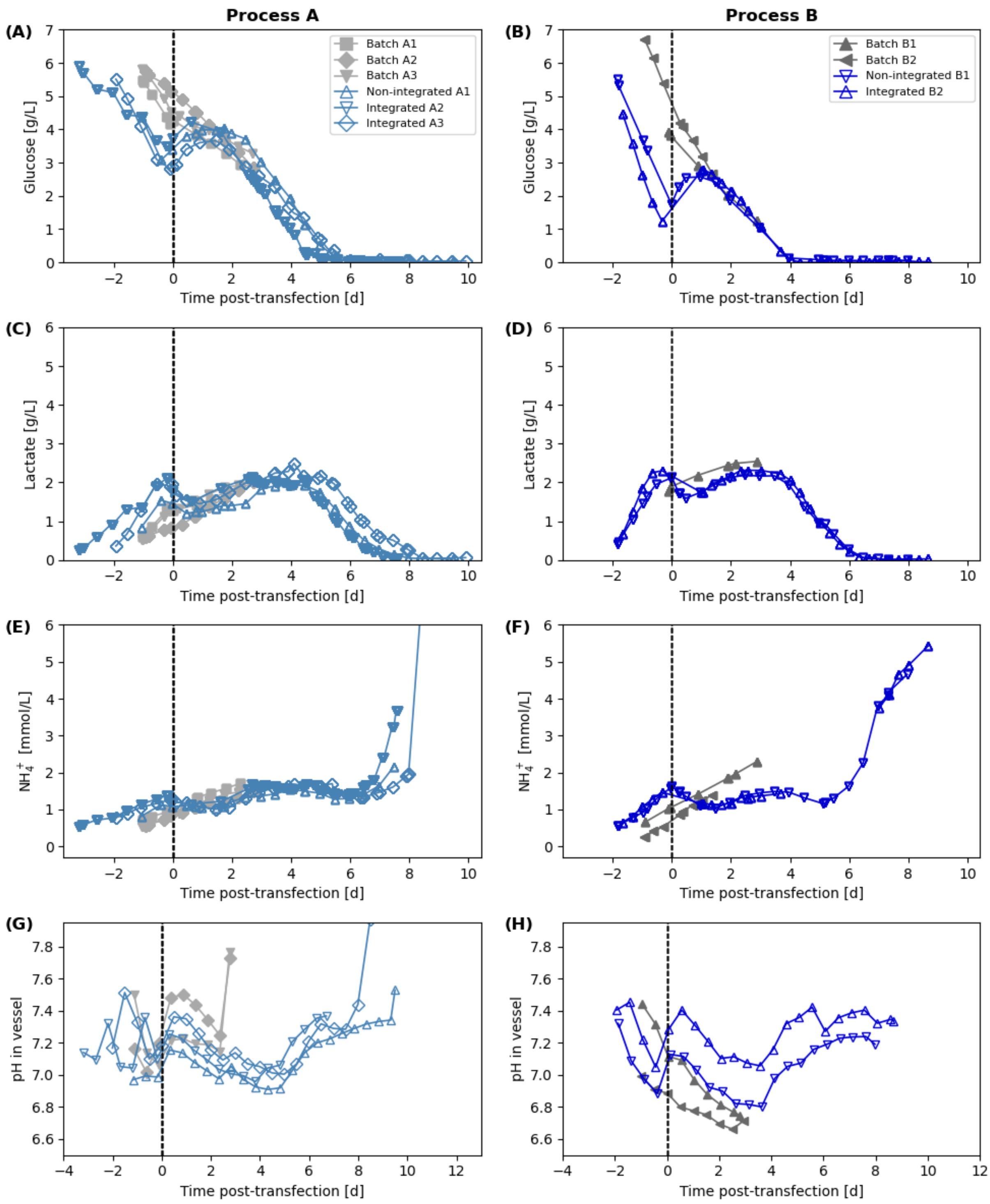


**Figure S2** Showing batch (grey-scale) and the continuous process (blue-scales). A and B visualizes the glucose levels, C and D for lactate, E and F ammonium, G and H the pH levels during the process times.

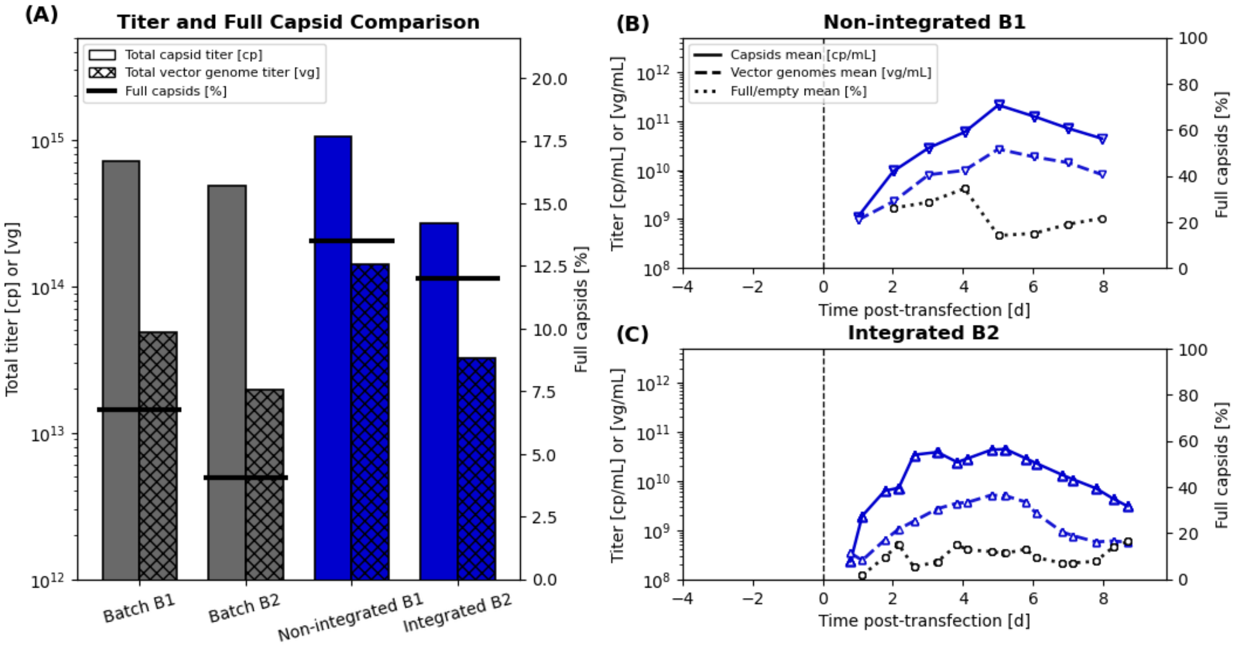


**Figure S3 Yield and Full/Empty Comparison.** A) Total yield of capsids [cp], vector genomes [vg] and full/empty ratio compared of the Process B Batch and Continuous. B) and C) shown as mean of the three measurement points: secreted positions from the vessel, after rTFF and after SF for capsids [cp/mL], of vector genomes [vg/mL] and the mean full/empty ratio.

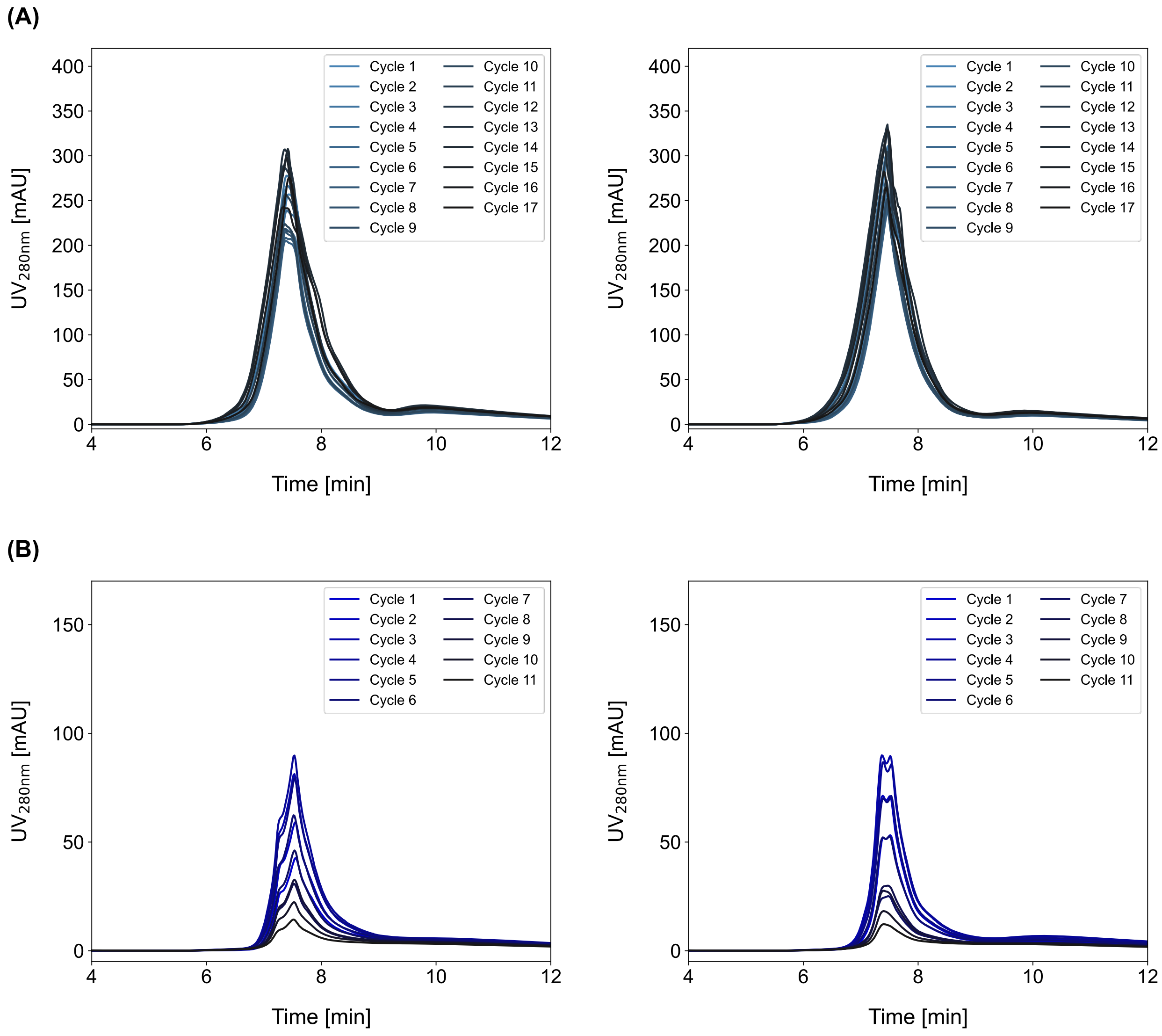


**Figure 4 Superimposed elution peaks of the integrated processes.** Elution peaks based on the $UV_{280nm}$ signal are shown for column 1 (left) and column 2 (right). Increasing cycle numbers are indicated by progressively darker colors.

**Table S1** Overview of downstream pocess parameters, comparing conventional batch operation and the integrated CaptureSMB. IC: Interconnected.

| Parameters | Units | Batch | Continuous |
|---|---|---|---|
| Equilibration length | [CV] | 10 | 10 |
| Equilibration | [cm/h] | 200 | 200 |
| (IC) Load flow rate | [cm/h] | 336 | 336 |
| (IC) PLW length | [CV] | 10 | 10 |
| (IC) PLW flow rate | [cm/h] | 100 | 100 |
| Parallel batch load flow rate | [cm/h] | N/A | variable |
| Elution length | [CV] | 10 | 10 |
| Elution flow rate | [cm/h] | 75 | 75 |
| Strip length | [CV] | 10 | 10 |
| Strip flow rate | [cm/h] | 200 | 200 |
| CIP length | [CV] | 25 | 25 |
| CIP flow rate | [cm/h] | 150 | 150 |
| Neutralization length | [CV] | 5 | 5 |
| Neutralization flow rate | [cm/h] | 200 | 200 |

**Table S2** Upstream process yields and space time yield (STY) of batch and continuous runs of the two processes.

| | | # | Yield [cp] | Yield [vg] | Space Time Yield (STY) [cp/L/d] | Space Time Yield (STY) [vg/L/d] |
|---|---|---|---|---|---|---|
| Process A | Batch (n=3) | 1 | $5.06 \times 10^{14}$ | N/A | $7.61 \times 10^{13}$ | N/A |
| | | 2 | $1.42 \times 10^{15}$ | N/A | $2.13 \times 10^{14}$ | N/A |
| | | 3 | $1.22 \times 10^{15}$ | N/A | $1.84 \times 10^{14}$ | N/A |
| | | mean | $1.05 \times 10^{15}$ | N/A | $1.58 \times 10^{14}$ | N/A |
| | | SD | $4.80 \times 10^{14}$ | N/A | $7.22 \times 10^{13}$ | N/A |
| | Continuous (n=3) | 1 | $3.99 \times 10^{15}$ | N/A | $2.71 \times 10^{14}$ | N/A |
| | | 2 | $3.81 \times 10^{15}$ | N/A | $2.30 \times 10^{14}$ | N/A |
| | | 3 | $4.39 \times 10^{15}$ | N/A | $2.48 \times 10^{14}$ | N/A |
| | | mean | $4.06 \times 10^{15}$ | N/A | $2.50 \times 10^{14}$ | N/A |
| | | SD | $2.95 \times 10^{14}$ | N/A | $2.04 \times 10^{13}$ | N/A |
| Process B | Batch (n=2) | 1 | $7.17 \times 10^{14}$ | $5.33 \times 10^{13}$ | $1.08 \times 10^{14}$ | $8.03 \times 10^{12}$ |
| | | 2 | $4.87 \times 10^{14}$ | $1.88 \times 10^{13}$ | $7.34 \times 10^{13}$ | $2.84 \times 10^{12}$ |
| | | mean | $6.02 \times 10^{14}$ | $3.61 \times 10^{13}$ | $9.07 \times 10^{13}$ | $5.43 \times 10^{12}$ |
| | Continuous (n=2) | 1 | $1.05 \times 10^{15}$ | $1.42 \times 10^{14}$ | $6.40 \times 10^{13}$ | $8.64 \times 10^{12}$ |
| | | 2 | $2.68 \times 10^{14}$ | $3.22 \times 10^{13}$ | $1.51 \times 10^{13}$ | $1.82 \times 10^{12}$ |
| | | mean | $6.61 \times 10^{14}$ | $8.73 \times 10^{13}$ | $3.95 \times 10^{13}$ | $5.23 \times 10^{12}$ |

**Table S3** Comparison of downstream performance between batch and integrated continuous operations for both process designs (A and B), with respect to load density, yield, productivity, and buffer consumption. For integrated operations, all calculations include the start-up and shutdown phases.

| | | Number of switches | Column volume | Load density | Yield | | Productivity | | Buffer consumption | |
|---|---|---|---|---|---|---|---|---|---|---|
| | | [#] | [mL] | [cp/mL resin] | [% cp] | [% vg] | [cp/L/h] | [vg/L/h] | [L/cp] | [L/vg] |
| Process A | Batch (n=2) | 1 | 0.491 | $2.19 \times 10^{16}$ | 87.4 | N/A | $9.05 \times 10^{15}$ | N/A | $4.17 \times 10^{-15}$ | N/A |
| | | 1 | 0.491 | $2.24 \times 10^{16}$ | 92.5 | N/A | $1.06 \times 10^{16}$ | N/A | $3.86 \times 10^{-15}$ | N/A |
| | Integrated (n=2) | 28 | 2×0.491 | $4.68 \times 10^{16}$ | 69.7 | N/A | $8.24 \times 10^{15}$ | N/A | $1.30 \times 10^{-15}$ | N/A |
| | | 32 | 2×0.491 | $6.26 \times 10^{16}$ | 66.4 | N/A | $1.24 \times 10^{16}$ | N/A | $1.01 \times 10^{-15}$ | N/A |
| Process B | Batch (n=2) | 1 | 0.491 | $2.36 \times 10^{13}$ | 82.2 | 85.0 | $9.36 \times 10^{15}$ | $5.21 \times 10^{14}$ | $4.13 \times 10^{-15}$ | $7.41 \times 10^{-14}$ |
| | | 1 | 0.491 | $2.36 \times 10^{13}$ | 85.6 | 65.9 | $9.76 \times 10^{15}$ | $4.04 \times 10^{14}$ | $3.96 \times 10^{-15}$ | $9.56 \times 10^{-14}$ |
| | Integrated (n=1) | 22 | 2×0.491 | $6.99 \times 10^{15}$ | 71.3 | 82.3 | $9.94 \times 10^{14}$ | $1.32 \times 10^{14}$ | $1.41 \times 10^{-14}$ | $1.07 \times 10^{-13}$ |
